# HMGB1/CXCL12 heterocomplex activates CXCR4 to suppress the ability of Müller glia to become proliferating progenitor-like cells in the chick retina

**DOI:** 10.64898/2026.11.09.765360

**Authors:** Lisa E. Kelly, Olivia B. Taylor, Heithem M. El-Hodiri, Samantha Lee, Elina Parekh, Evan C. Hawthorn, Joshua Wright, Mary R. Guest, Allen Tangeman, Andy J. Fischer

## Abstract

We investigated how High Mobility Group Box 1 (HMGB1) signaling regulates glial reactivity and Müller glia-derived progenitor cell (MGPC) formation in the chick retina. HMGB1 released from stressed cells is known to form a heterocomplex with C-X-C chemokine ligand 12 (CXCL12) to activate C-X-C chemokine receptor 4 (CXCR4). HMGB1 is broadly expressed in retinal neurons and glia and is rapidly exported from the nuclei of stressed inner retinal neurons after damage, a process blocked by inflachromene. Stressed or damaged amacrine cells also exported different transcription factors out nuclei. *CXCL12* is expressed by amacrine, horizontal, and bipolar cells, whereas *CXCR4* is expressed by Müller glia and is rapidly upregulated following damage. CXCR4 agonist or recombinant HMGB1/CXCL12 treatment selectively stimulates mTOR and Jak/Stat signaling in Müller glia in undamaged retinas. Inhibition of CXCR4 signaling, HMGB1 nuclear export, or HMGB1/CXCL12 complex formation potently stimulated MGPC proliferation without affecting neuronal death or microglial activation. In retinas treated with insulin and FGF2, to induce MGPC proliferation independent of injury, HMGB1 remains confined to nuclei, *CXCR4* is downregulated, and inhibition of HMGB1 or CXCR4 has no effect. We further demonstrate that microglia are required to maintain *CXCR4* expression in Müller glia, and blockade of HMGB1/CXCL12-CXCR4 signaling does not restore MGPC proliferation in microglia-depleted retinas. Together, these findings identify HMGB1/CXCL12-CXCR4 signaling as an injury-induced pathway that limits Müller glia reprogramming and MGPC formation in damaged retinas.

## Introduction

Different cell signaling pathways regulate retinal inflammation <u>and</u> impact the reprogramming of Müller glia (MG) into MG-derived progenitor cells (MGPCs) in the retinas of different vertebrate species (Taylor et al., 2025b). Nuclear Factor Kappa B (NFκB) is a master regulator of injury responses in MG, acts to restore glial quiescence and suppress the neurogenic capacity of MG-derived progenitors in mice (Hoang et al., 2020; Palazzo et al., 2020). NFκB-signaling in MG in damaged retinas is driven by pro-inflammatory signals produced by activated microglia (Palazzo et al., 2022, 2023). Inhibition of NFκB-signaling reduces immune cell recruitment in damaged retinas, enhances neuronal survival, and increases the generation of neuron-like cells from Ascl1-overexpressing MG (Palazzo et al., 2022). In the chick retina, by comparison, rapid activation of NFκB by microglia-derived signals initiates the process of MG activation and reprogramming, whereas sustained NFκB signaling suppresses MGPC proliferation and neuronal differentiation of progeny (Palazzo et al., 2020).

High Mobility Group Box 1 (HMGB1) protein is a highly conserved protein that is widely expressed by neuronal cells from embryonic stages through adulthood (Tang et al., 2023). Although HMGB1 mediates intracellular functions, it can also be secreted to elicit extracellular functions in development, disease, and inflammation (Fig. 1). HMGB1 is released into the extracellular space through the secretory lysosomal pathway or by passive release from dying or stressed cells (Fig. 1; reviewed by (Chen et al., 2022)). Active secretion occurs in response to cytokines, cellular stress and activation of signaling pathways including NFkB, Notch, MAPK and STAT (Chen et al., 2022). The secretion of HMGB1 is controlled by post-translational modifications including N-glycosylation at N37, N134 and N135, as well as post-translational modifications, including acetylation, ADP-ribosylation, phosphorylation, and methylation (reviewed by (Chen et al., 2022). Extracellular HMGB1 can regulate injury responses by binding directly to toll-like receptors 2 and 4 (TLR2 and TLR4), or forming a heterocomplex with C-X-C chemokine ligand 12 (CXCL12) to activate C-X-C chemokine receptor 4 (CXCR4; Fig. 1). CXCR4 is expressed by different types of cells, including hematopoietic and endothelial cells (ECs), neurons and stem cells (Lee et al., 2018; Bianchi and Mezzapelle, 2020). Following injury, HMGB1/CXCL12 heterocomplex activates CXCR4 to influence skeletal, hematopoietic, and muscle regeneration *in vivo* (Bianchi and Mezzapelle, 2020). HMGB1 signaling through TLR4 or CXCL12-CXCR4 has been implicated in the pathology of diabetic retinopathy and the pathogenesis of glaucoma and age-relate macular degeneration (Liu et al., 2018). However, the role of HMGB1/CXCL12-CXCR4 in mediating communication between damaged neurons, microglia and MG – and its involvement in MG reprogramming into neurogenic MGPCs – remains unknown.

**Figure 1:**
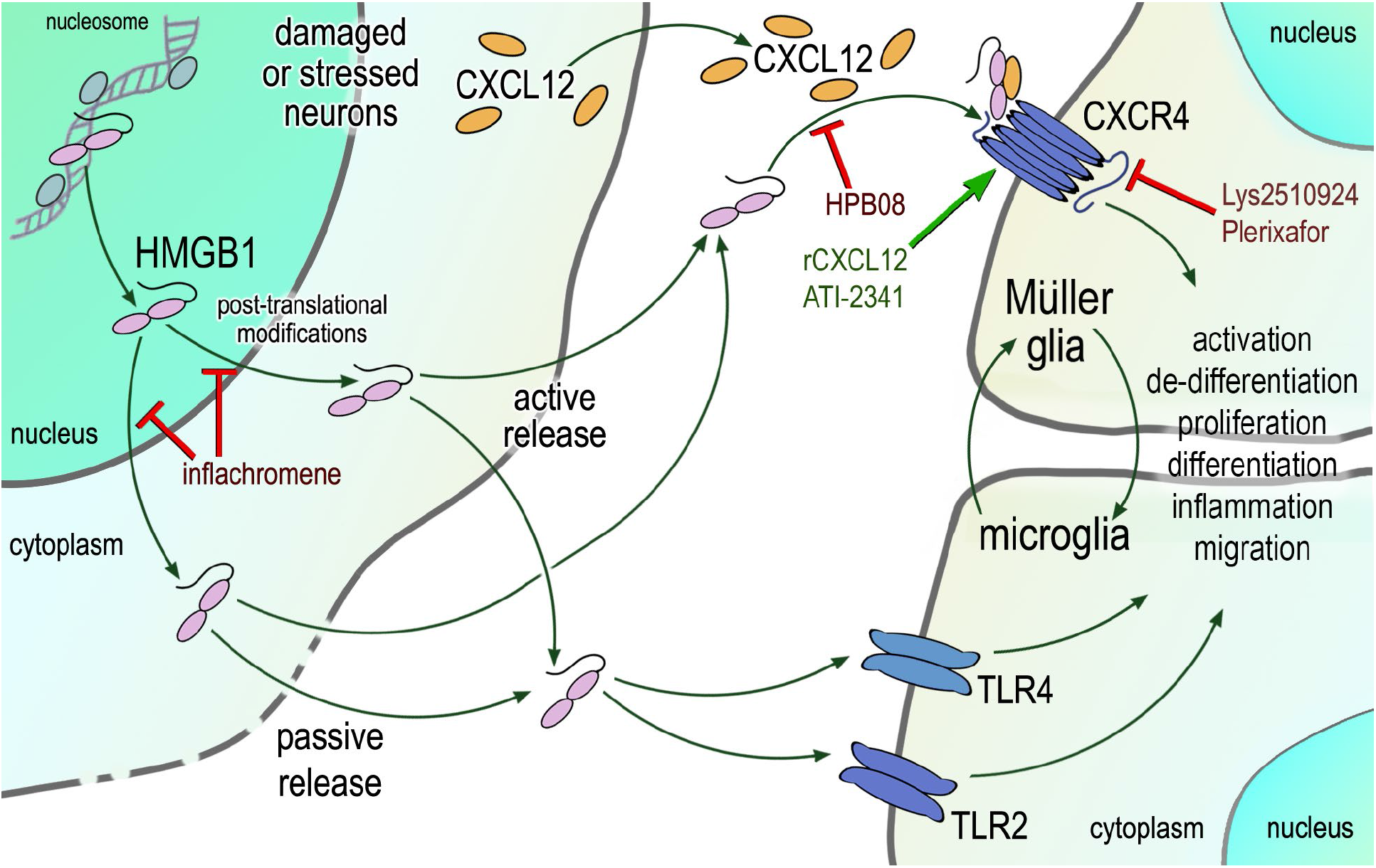
Schematic summary of HMGB1 signaling in the damaged retina. Antagonists are indicated by red text and lines and agonists are indicated by green text and arrows.

Recent evidence indicates that the key “cellular decisions” to become reactive and then return to quiescence *<u>or</u>* transition to a progenitor-like state occur rapidly, within 3-12 hours after neuronal injury (Hoang et al., 2020; Taylor et al., 2025b). Currently, the signals that initiate and coordinate the responses of microglia and MG to neuronal damage are poorly understood. The identity of the damage-specific factors that impact MG activation and reprogramming into MGPCs remains poorly defined. The purpose of this study was to investigate whether HMGB1 signaling is involved in the formation of MGPCs in the chick retina.

## Methods

### Animals

The animal use approved in these experiments was in accordance with the guidelines established by the National Institutes of Health and IACUC at The Ohio State University. Newly hatched P0 wildtype leghorn chicks (*Gallus gallus domesticus*) were obtained from Meyer Hatchery (Polk, Ohio). Post-hatch chicks were maintained in a regular diurnal cycle of 12 hours light, 12 hours dark (8:00 AM-8:00 PM). Chicks were housed in stainless-steel brooders at 25°C and received water and Purina^TM^ chick starter *ad libitum*.

### Intraocular Injections

Chicks were anesthetized with 2.5% isoflurane mixed with oxygen from a non-rebreathing vaporizer. The technical procedures for intraocular injections were performed as previously described (Fischer et al., 1998, 2005). With all injection paradigms, pharmacological and vehicle treatments were administered to the right and left eye respectively. Compounds were injected in 20 μl sterile saline. Hydrophobic compounds were injected with <30% DMSO in saline, according to solubility data provided by the vendor. 5-Ethynyl-2’-deoxyuridine (EdU) was injected into the vitreous chamber to label proliferating cells. Compounds included in this study are described in **Table 1**. For chicks between P7 and P14, we estimated volumes of 100 μl of liquid vitreous and 800 μl gel vitreous, and an average eye weight of 0.9 grams (Oishi and Murakami, 1985; Prashar et al., 2009). Table 1 includes the ranges of reported *in vivo* ED50 (50% effective dose; mg/kg) and the calculated initial maximum dose (mg/kg equivalent per eye). Doses were chosen based on estimates of the initial maximum ocular dose that were within the range of reported ED50. Injection paradigms are included in each figure.

**Table 1.**
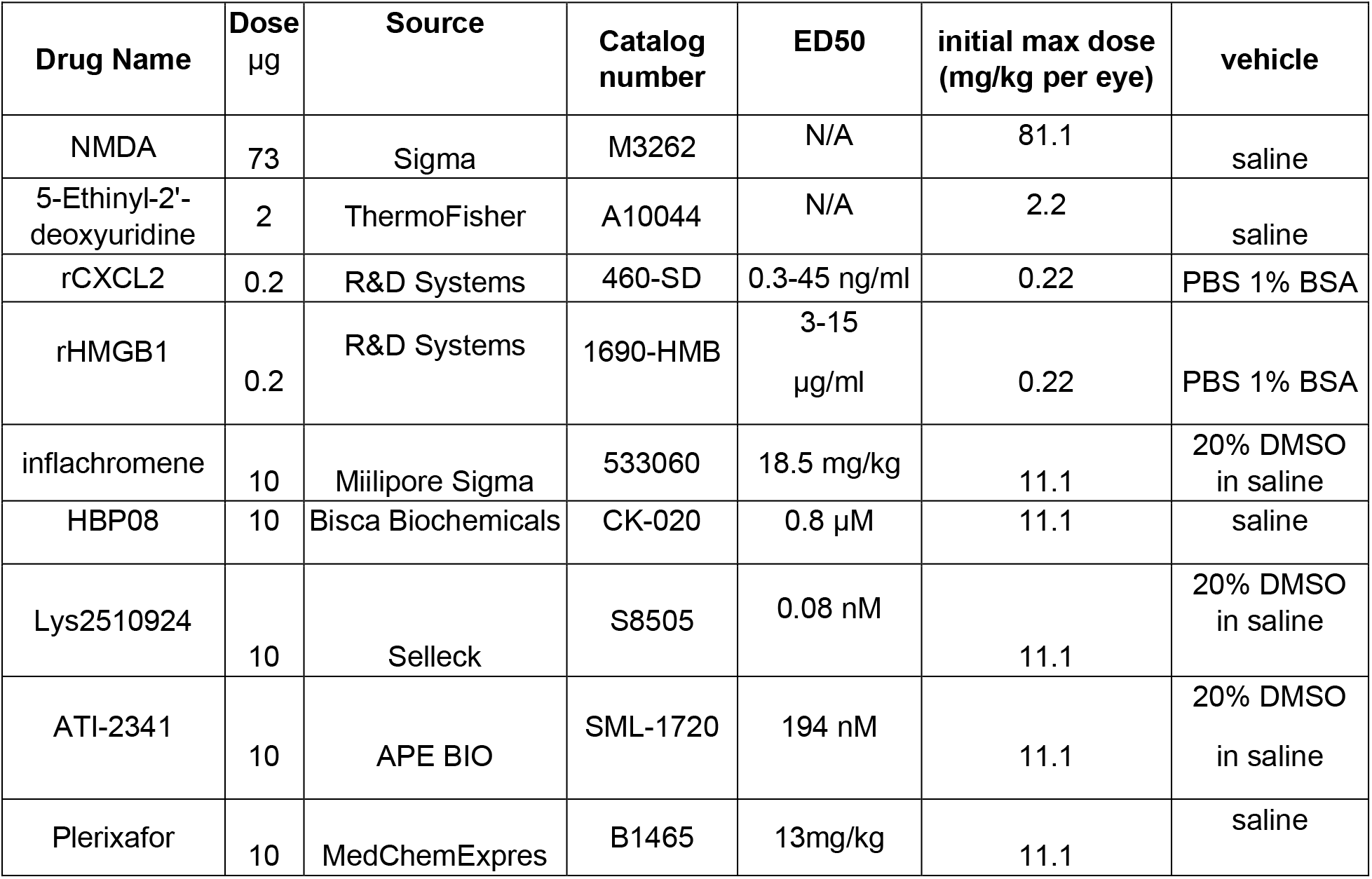
Pharmacological Compounds.

| **Drug Name** | **Dose**<br>µg | **Source** | **Catalog number** | **ED50** | **initial max dose (mg/kg per eye)** | **vehicle** |
|---|---|---|---|---|---|---|
| NMDA | 73 | Sigma | M3262 | N/A | 81.1 | saline |
| 5-Ethinyl-2'-deoxyuridine | 2 | ThermoFisher | A10044 | N/A | 2.2 | saline |
| rCXCL2 | 0.2 | R&D Systems | 460-SD | 0.3-45 ng/ml | 0.22 | PBS 1% BSA |
| rHMGB1 | 0.2 | R&D Systems | 1690-HMB | 3-15<br>µg/ml | 0.22 | PBS 1% BSA |
| inflachromene | 10 | Miilipore Sigma | 533060 | 18.5 mg/kg | 11.1 | 20% DMSO in saline |
| HBP08 | 10 | Bisca Biochemicals | CK-020 | 0.8 µM | 11.1 | saline |
| Lys2510924 | 10 | Selleck | S8505 | 0.08 nM | 11.1 | 20% DMSO in saline |
| ATI-2341 | 10 | APE BIO | SML-1720 | 194 nM | 11.1 | 20% DMSO<br>in saline |
| Plerixafor | 10 | MedChemExpres | B1465 | 13mg/kg | 11.1 | saline |

### Preparation of clodronate liposomes

Clodronate liposomes were prepared as described previously (Van Rooijen, 1989; van Rooijen, 1992; Van Rooijen and Sanders, 1994; Fischer et al., 2014). In short, 50 ng cholesterol (Sigma C3045) and 8 mg L-α-phosphatidyl-DL-glycerol sodium salt (Sigma P8318) were dissolved in chloroform and evaporated into a thin film in a round-bottom flask. Then, the liposome film was dissolved in sterile PBS containing 158 mg dichloro-methylene diphosphonate (Sigma D4434). The clodronate was encapsulated by the liposomes via sonication at 42,000 Hz. The clodronate liposomes in PBS were centrifuged at 10,000 RCF, gently resuspended in 150 ml sterile PBS, and injected 20 ul/eye immediately. Our previous studies have shown that this method ablates >95% of microglia/macrophages 2-3 days after intraocular injection (Zelinka et al., 2012; Fischer et al., 2014; El-Hodiri et al., 2023).

### Fixation, sectioning, and immunocytochemistry

Retinas were fixed, sectioned, and immunolabeled as described previously (Fischer et al., 1998, 2005; Campbell et al., 2019). To identify MG, we labeled sections for Sox2 or Sox9. Antibodies to Sox2 and Sox9 are known to label the nuclei or MG in the INL and the nuclei of astrocytes at the vitread surface of the retina (Fischer et al., 2010). None of the observed labeling was due to non-specific labeling of secondary antibodies or auto-fluorescence because sections labeled with secondary antibodies alone were devoid of fluorescence. Primary antibodies used in this study are described in **Table 2**. Secondary antibodies included donkey-anti-goat-Alexa488/594/647 (Life Technologies A11055; A11058; A21447), donkey-anti-rabbit-Alexa488/594 (Life Technologies A21206; A21207); and goat-anti-mouse-Alexa488/568 (Life Technologies A11001; A-11004) diluted to 1:1000 in PBS plus 0.2% Triton X-100. Nuclear staining was accomplished using DAPI (MilliporeSigma 5.08741.0001).

**Table 2:**
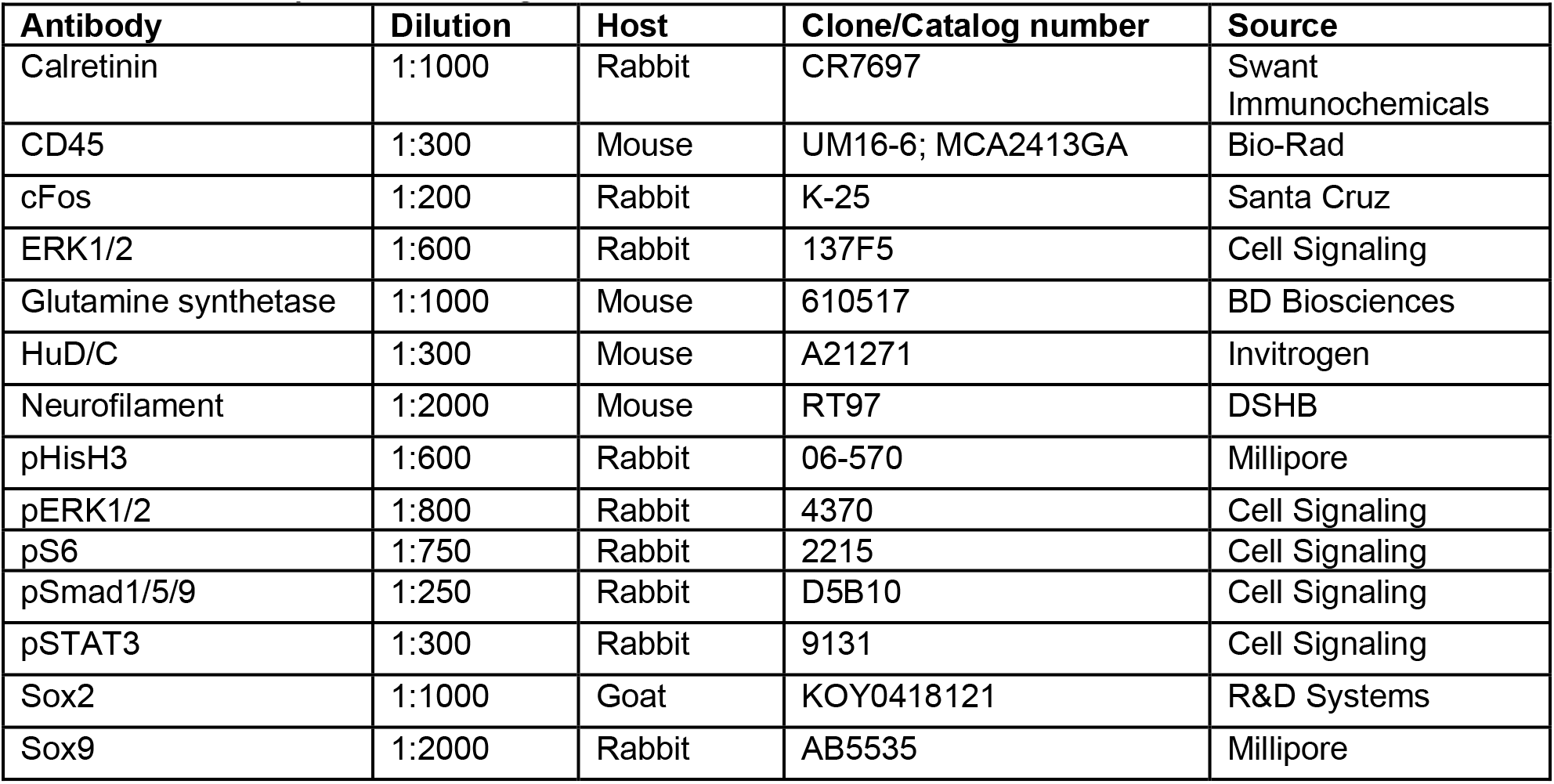
Antibody table – antigen, dilution, host and sources of antibodies.

### Labeling for 5-Ethynyl-2-deoxyuridine (EdU)

For the detection of nuclei that incorporated EdU, immunolabeled sections were fixed in 4% formaldehyde in 0.1M PBS pH 7.4 for 5 minutes at room temperature. Samples were washed for 5 minutes with PBS, permeabilized with 0.5% Triton X-100 in PBS for 1 minute at room temperature and washed twice for 5 minutes in PBS. Sections were incubated for 30 minutes at room temperature in a buffer consisting of 100 mM Tris, 8 mM CuSO4, and 100 mM ascorbic acid in dH2O. The Alexa Fluor 647 Azide (Thermo Fisher Scientific A10277) was added to the buffer at a dilution of 1:500.

### Terminal deoxynucleotidyl transferase dUTP nick end labeling (TUNEL)

The TUNEL method was used to identify dying cells with fragmented DNA. We used an *in-situ* Cell Death Detection kit from Roche (Fluorescein, 11684795910) according to manufacturer instructions.

### Photography, immunofluorescence measurements, and statistics

Wide-field photomicroscopy was performed using a Leica DM5000B microscope equipped with epifluorescence and Leica DC500 digital camera. Confocal images were obtained using a Leica SP8 imaging system at the Department of Neuroscience Main Campus Imaging Facility at the Ohio State University. Images were optimized for color, brightness and contrast, multiple channels over-laid and figures constructed using Adobe Photoshop. Cell counts were performed on representative images. To avoid the possibility of region-specific differences within the retina, cell counts were consistently made from the same region of retina for each data set. Peripheral regions of the retina were designated as 1 to 3 mm from the circumferential marginal zone, and central regions of the retina were designated as within 3 mm of the posterior pole of the eye in the nasotemporal plane.

Similar to previous reports (Ghai et al., 2009; Gallina et al., 2015b; Taylor et al., 2025a), immunofluorescence was quantified by using Image J (NIH). Identical illumination, microscope and camera settings were used to obtain images for quantification. Measurements of pS6, pStat3 and cFos immunofluorescence in MG were made for a fixed, cropped area of INL or INL+ONL. Measurements in the nuclei of MG were made by selecting the total area of pixels above threshold for Sox2 (in the red channel), copying pStat3 or cFos (in the green channel) into new images and measuring pixels intensities. Measurements were made for regions containing pixels with intensity values of 70 or greater (0 = black and 255 = saturated). The mean intensity was calculated as the mean pixel values for all pixels within threshold regions.

Significance of difference between two treatment groups accounting for inter-individual variability (means of treated-control values) was determined using a two-tailed, paired t-test. Where significance of difference was determined between two treatment groups, we performed a two-tailed, paired or unpaired t-test. Significance of difference between multiple groups was determined using one-way ANOVA followed by and Šidák’s test for multiple comparisons. GraphPad Prism 11 was used for statistical analyses and generation of histograms and bar graphs.

### scRNA-seq

We analyzed scRNA-seq libraries that were generated and characterized previously (Hoang et al., 2020; Campbell et al., 2021b; El-Hodiri et al., 2023). Dissociated cells were loaded onto the 10X Chromium Cell Controller with Chromium 3’ V2, V3 or Next GEM reagents. Using Seurat toolkits (Powers and Satija, 2015; Satija et al., 2015), Uniform Manifold Approximation and Projection (UMAP) for dimensional reduction plots were generated from 9 separate cDNA libraries, including 2 replicates of control undamaged retinas, and retinas at different times after NMDA-treatment. Seurat was used to construct gene lists for differentially expressed genes (DEGs), violin/scatter plots, and dot plots. Significance of difference in violin/scatter plots was determined using a Wilcoxon Rank Sum test with Bonferroni correction. Genes that were used to identify different types of retinal cells included the following: (1) Müller glia: *GLUL, VIM, SCL1A3, RLBP1*, (2) MGPCs: *PCNA, CDK1, TOP2A, ASCL1*, (3) microglia: *C1QA, C1QB, CCL4, CSF1R, TMEM22*, (4) ganglion cells: *THY1, POU4F2, RBPMS2, NEFL, NEFM*, (5) amacrine cells: *GAD67, CALB2, TFAP2A*, (6) horizontal cells: *PROX1, CALB2, NTRK1*, (7) bipolar cells: *VSX1, OTX2, GRIK1, GABRA1*, and (7) cone photoreceptors: *CALB1, GNAT2, GNB3, OPN1LW*, and (8) rod photoreceptors: *RHO, NR2E3, ARR3*.

## Results

### Expression of CXCR4 and HMGB’s in the developing chick retina

We first analyzed patterns of expression in the embryonic chick retina using previously generated single cell RNA-sequencing (scRNA-seq) datasets (Campbell et al., 2021a). After filtering to remove cells with low gene counts, doublets, and high mitochondrial content, the combined dataset comprised 22,698 cells from E5, E8, E12, and E15 retinas (Supplemental Fig. 1a). UMAP embedding of aggregated libraries revealed distinct clusters of cells corresponding to developmental stage and cell type (Supplemental Fig. 1a,b). Embryonic retinal progenitor cells (eRPCs) from E5 and E8 retinas were identified by expression of *ASCL1, CDK1*, and *TOP2A* (Supplemental Fig. 1c). Immature Müller glia (iMG) were identified by increasing expression of *NOTCH1, HES5*, and *FABP7* (not shown), whereas maturing Müller glia (mMG) exhibited elevated expression of *GLUL, RLBP1*, and *SLC1A3* (Supplemental Fig. 1d). *HMGB1, HMGB2*, and *HMGB3* were highly expressed in eRPCs from E5 and E8 retinas but were significantly downregulated in iMG at E8 and in mMG at E12 and E15 (Supplemental Fig. 1e,f). *HMGB1* and *HMGB3* were broadly expressed across all types of differentiating retinal neurons (Supplemental Fig. 1e). In contrast, *CXCR4* expression was low and restricted to a small subset of eRPCs at E5 (Supplemental Fig. 1e,f), but was significantly upregulated in iMG at E8 and in mMG at E12 and E15 (Supplemental Fig. 1e,f). These findings suggest that the maturation of MG involves the downregulation of *HMGB1/2/3* and signaling through *CXCR4.* Further, these findings predict that de-differentiation and reprogramming of MG into proliferating MGPCs might involve upregulation of *HMGB1/2/3*.

### Expression of *CXCR4, CXCL12* and *HMGB*’s in damaged chick retinas

We next probed for patterns of expression in normal and damaged retinas from hatched chicks. We analyzed aggregated scRNA-seq datasets of >180,000 cells isolated from retinas treated with saline, NMDA, 2 or 3 doses insulin+FGF2 or the combination of NMDA and insulin+FGF2 (Fig. 2a). We have previously used these scRNA-seq libraries to characterize expression patterns of genes related to cell-signaling and gene networks (Campbell et al., 2021b; El-Hodiri et al., 2022; Palazzo et al., 2022; Taylor et al., 2024, 2025a)

**Figure 2:**
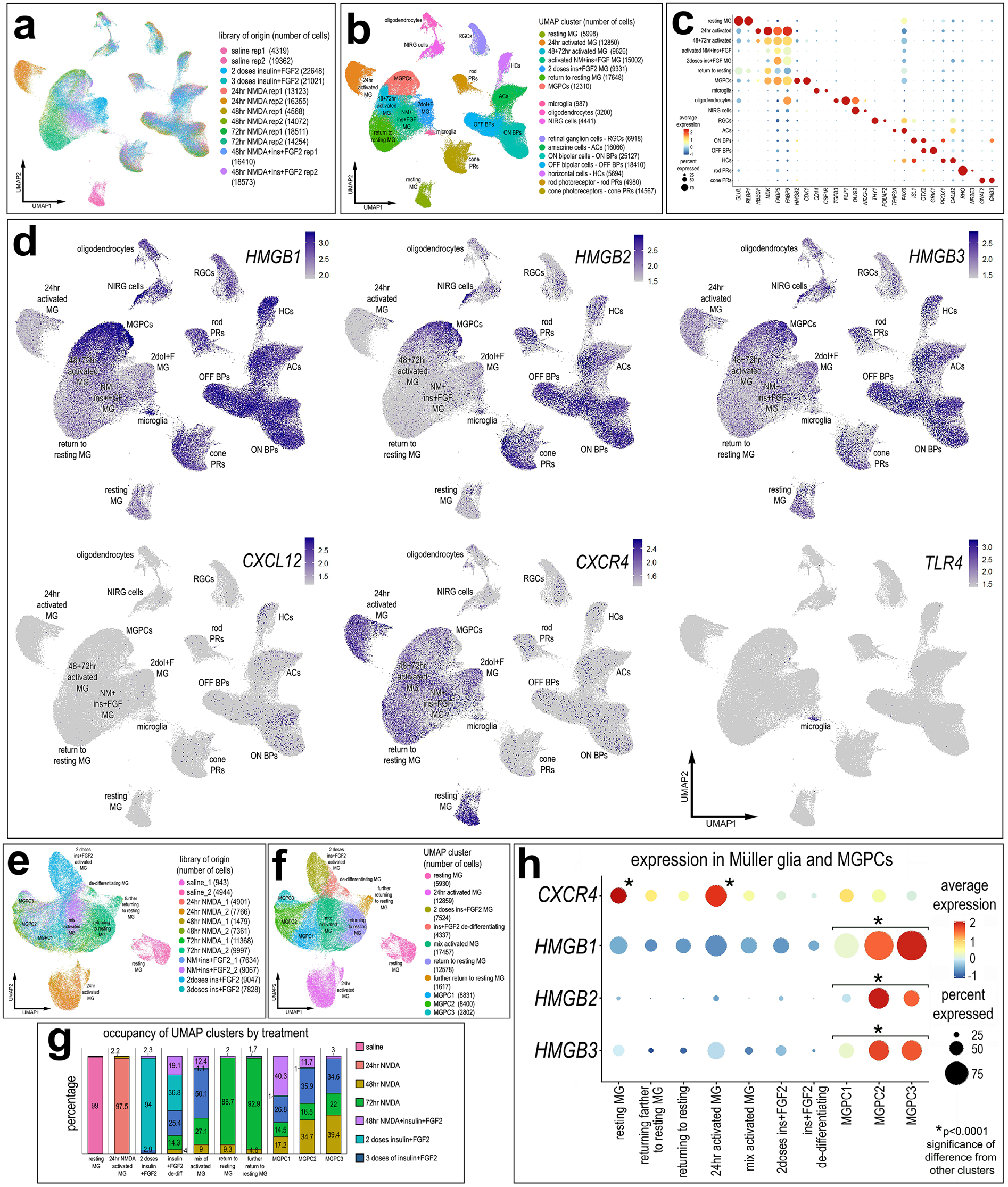
Patterns of expression of genes in resting MG, activated MG, and MGPCs. These scRNA-libraries were generated and analyzed previously (Hoang et al., 2020; Palazzo et al., 2020; Campbell et al., 2021). Aggregate scRNA-seq libraries were generated for cells from control retinas and retinas 24, 48, and 72 h after NMDA-treatment, retinas treated with two or three doses of insulin and FGF2, and retinas treated with insulin, FGF2 and NMDA (**a**). Clusters of cells were identified based on distinct patterns of gene expression (**b**). The dotplot in **c** illustrates the levels (average expression heatmap) and percentage of expressing cells (dot size) for representative genes that distinguish different clusters and types of neurons and glia. UMAP feature plots illustrate patterns and levels of expression of *HMGB1, HMGB2, HMGB3, CXCL12, CXCR4* and *TLR4* (**d**). MG were bioinformatically isolated and re-embedded in a UMAP plot for further analyses (**e-h**). UMAP-ordered cells formed distinct clusters of resting MG, returning to resting MG, 24hr activated MG, activated MG and MGPCs based on distinct patterns of gene expression that were correlated to occupancy by treatment (**g**). The dotplot in **h** illustrates the levels (average expression heatmap) and percentage of expressing cells (dot size) for *CXCR4*, *HMGB1, HMGB2* and *HMGB3* in different clusters of MG and MGPCs. Significance of difference (*p<<0.0001) was determined using Wilcox rank sum with Bonferoni correction (see supplemental Table 1 for statistics). Abbreviations: MG - Müller glia; NMDA - N-methyl-D-aspartate; UMAP - uniform manifold approximation and projection.

UMAP plots revealed clusters of different retinal cell types that were identified based on well-established markers (Fig. 2b,c), as described in the methods. *HMGB1/2/3* were widely expressed by all types of retinal neurons and glia, and *CXCL12* was expressed by scattered bipolar, amacrine and horizontal cells (Fig. 2d). *CXCR4* was expressed predominantly by resting MG, activated MG and MGPCs (Fig. 2d). By comparison, *TLR4* was detected only in microglia (Fig. 2d); insufficient numbers of cells precluded comparisons across timepoints after damage. We isolated and re-embedded the MG from different treatment groups into a UMAP plot and determined the percent occupancy of MG from different treatments into different clusters (Fig. 2e-g). Resting MG, activated MG from 24 hrs after NMDA-treatment and MG treated with two doses of insulin+FGF2 formed distinct clusters of cells (Fig. 2e-g). MGPCs formed discrete regions of cells wherein cell cycle progression correlated with spatial segregation in the UMAP plot (Fig. 2e-g). We found that *CXCR4* expression was highest in resting MG and 24hr activated MG, whereas *CXCR4* was significantly downregulated in MG and MGPCs from other treatment groups (Fig. 2h). By comparison, levels of *HMGB1*, *HMGB2* and *HMGB3* were highest in MGPCs (Fig. 2h).

We next analyzed scRNA-seq libraries that were aggregated from control retinas and retinas treated with NMDA-damage at early times (3, 12, and 48 hrs) after treatment, as described previously (Campbell et al., 2021a). Consistent with the other scRNA-seq libraries, *HMGB1* and *HMGB2* were widely expressed by different cells, including INL neurons and retinal ganglion cells (RGCs; Fig. 3c,e). Similarly, *CXCL12* was expressed by inner retinal neurons, including amacrine, horizontal and bipolar cells (Fig. 3c,e). Within 3 hrs of NMDA-treatment, *HMGB1*, *HMGB2* and *CXCL12* were rapidly downregulated in whole retina, specifically in microglia, INL neurons and RGCs (Fig. 3e). *CXCL12* remained at low levels, whereas *HMGB1* and *HMGB2* were upregulated by 48hrs after treatment (Fig. 3e). *CXCR4* was exclusively expressed by MG and was highly upregulated from 3 to 12hrs and then downregulated by 48hrs after NMDA-treatment (Fig. 3c,d). Consistent with Figure 2, *TLR2* and *TLR4* were expressed by only microglia (Fig. 3c,e). Secreted HMGB1 is known to act at SIGLEC-10, TLR9 and RAGE (*AGER*) receptors (Chen et al., 2022). However, *SIGLEC10*, *TLR9* and *AGER* were not detected in scRNA-seq libraries of chick retinas (data not shown).

**Figure 3.**
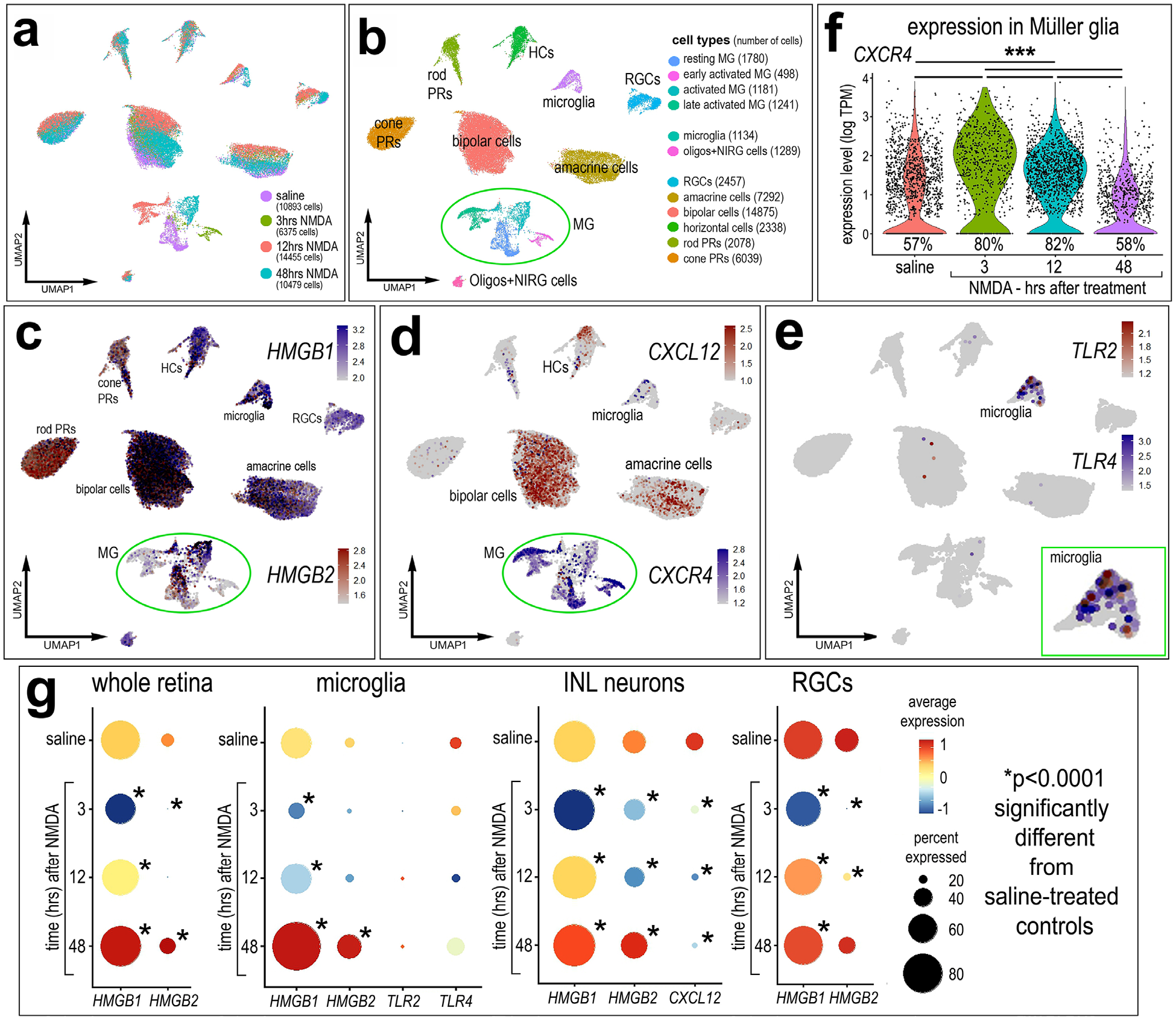
Patterns of expression of *HMGB1, HMGB2, CXCL12, CXCR4, TLR2 and TLR4* are rapid and dynamic in damaged chick retinas. These scRNA-libraries were generated and analyzed previously (Campbell et al., 2021b eCB paper). scRNA-seq libraries were generated from cells obtained from undamaged retinas and from retinas damaged by NMDA at 3, 12 and 48 hrs after treatment (**a**). Clusters of MG were identified based on expression of resting and activated markers such as *GLUL* and *TGFB2* (**b**). Feature plots in **c, d** and **e** illustrate patterns and levels of expression of *HMGB1, HMGB2, CXCL12, TLR2* and *TLR4* in UMAP clustered retinal cells. The violin plot in **f** illustrates levels and percent expression in MG in control retinas and at different times after NMDA-treatment. Dotplots in **g** illustrate the average expression (heatmap) and percentage of expressing cells (dot size) for *HMGB1, HMGB2, TLR2, TLR4* and/or *CXCL12* in whole retina, microglia, INL neurons or RGCs. Wilcoxon Rank Sum Test with Bonferroni correction was used to determine the significance of difference (*p<0.0001, ***p<<0.0001). Abbreviations: MG – Muller glia; UMAP, uniform manifold approximation and projection; INL – inner nuclear later; RGCs – retinal ganglion cells; HCs – horizontal cells; PRs – photoreceptor cells.

### Immunolabeling for HMGB1

Expression patterns of HMGB1 were validated using immunofluorescence. Consistent with data from scRNA-seq libraries, HMGB1 protein was widely detected in the nuclei of nearly all cells in undamaged, normal retinas (Fig. 4a; supplemental Fig.2). We detected relatively few cells that had little or no immunoreactivity for HMGB1 in the nuclei or cytoplasm; these cells were found at the borders of the IPL and ONL/INL and the IPL and INL/GCL (Fig. 4a). The HMGB1-negative cells had normal patterns of DAPI fluorescence filling out the nuclei (Fig. 4a). HMGB1 was detected in all Sox2-positive MG nuclei, and the nuclei of horizontal, amacrine and ganglion that were labeled for calretinin (Supplemental Fig. 2). In the GCL, we often observed large HMGB1 puncta in the cytoplasm of RGCs (Fig. 4b, 5b, Supplemental Fig. 2).

**Figure 4:**
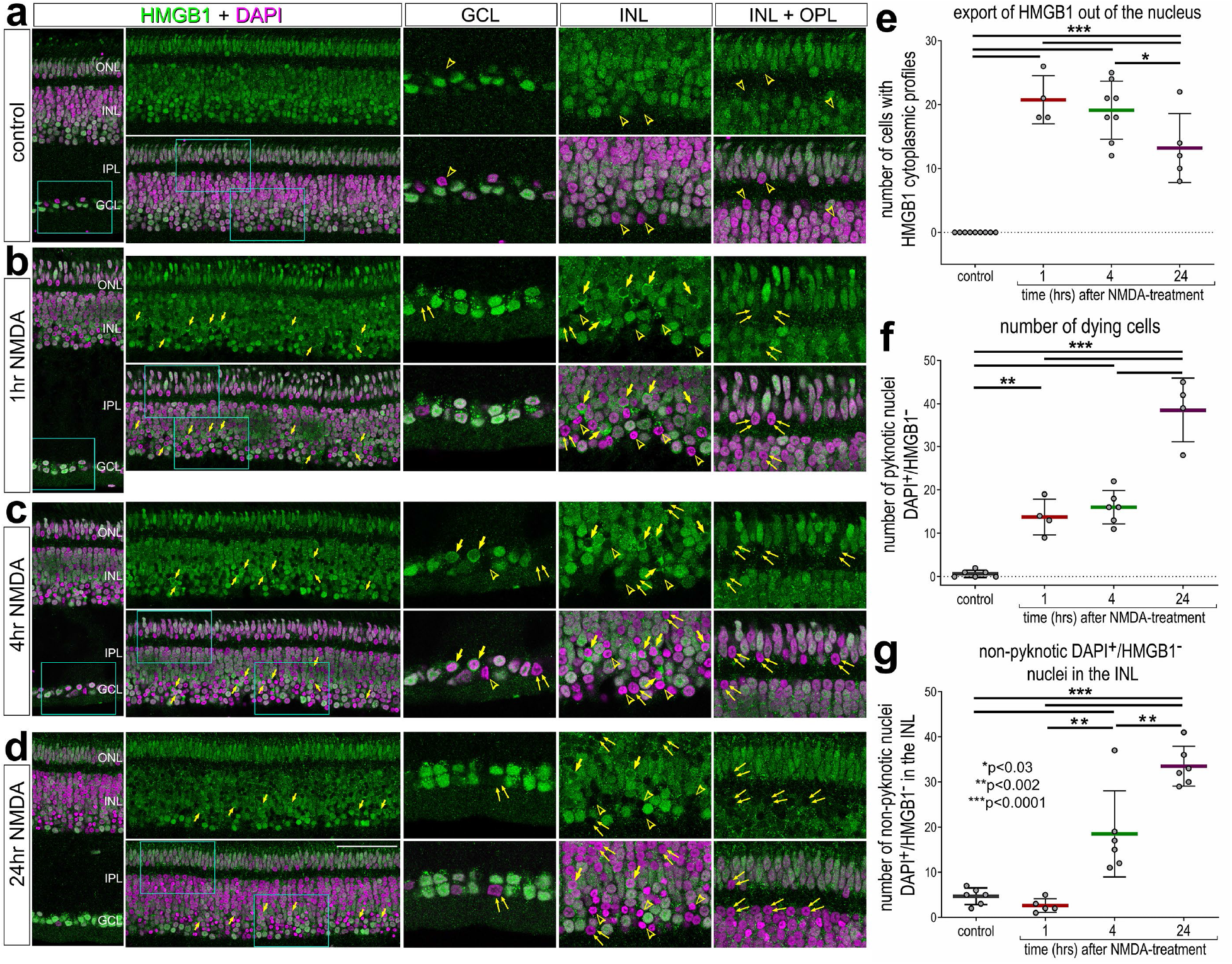
HMGB1 is rapidly exported out of the nuclei of stressed neurons. Eyes were injected with saline or NMDA and retinas harvested at 1, 4 or 24 hrs later. Retinal sections were labeled with DAPI (magenta) and antibodies to HMGB1 (green; **a-d**). Confocal microscopy as used to obtain single optical sections that were less than 2 µm thick. Hollow arrowheads (yellow) indicate DAPI-positive pyknotic nuclei that are negative for HMGB1. Arrows (yellow) indicate DAPI-intense nuclei with HMGB1 exported out of the nuclei into the peri-nuclear cytoplasm. Small double-arrows (yellow) indicate non-pyknotic DAPI-positive cells that have little or no HMGB1-immunoreactivity. The calibration bars in **d** represent 50 µm and applies to **a-d**. Areas of the ONL, INL and GCL that are indicated by cyan rectangles are enlarged 2-fold in adjacent panels. Histograms (**e**,**f,g**) illustrate the mean (bar ± SD), each dot represents one biological replicate. Significance of difference (p-values) was determined by one-way ANOVA and Tukey’s multiple comparison test. Abbreviations: ONL – outer nuclear layer, INL – inner nuclear layer, IPL – inner plexiform layer, GCL – ganglion cell layer, ns – not significant. NMDA, N-methyl-D-aspartate.

At 1 and 4 hrs after NMDA-treatment, we detected numerous putative pyknotic nuclei in the inner INL and GCL (Fig. 4b,c,f). These pyknotic nuclei contained condensed chromatin that was intensely labeled for DAPI (Fig. 4b,c,f). Pyknotic nuclei were most abundant at 24 hrs after NMDA (Fig. 4f) which is consistent with the time-course of abundance for TUNEL-positive cells (Fischer et al., 1998, 2004; Gallina et al., 2015b). Many of these pyknotic nuclei had HMGB1-immunoreactivity in the cytoplasm with HMGB1 diminished or absent from the nucleus; these cells were found predominantly in the amacrine cell layer of the INL (Fig. 4b,c,e). At 24hrs after NMDA-treatment, we detected increased numbers of pyknotic nuclei in the INL (Fig. 4d,f). These DAPI-intense cells were consistently negative for HMGB1 in nucleus, and relatively few of these cells had HMGB1 in the peri-nuclear cytoplasm (Fig. 4d,e). Pyknotic nuclei (intense DAPI-labeling and irregular shape) in the INL with HMGB1 in the cytoplasm were significantly less abundant at 24hrs than at 1 or 4 hrs after NMDA (Fig. 4d). We also identified non-pyknotic cells with HGMB1 diminished or absent from the nucleus. We hypothesized that these cells may have been stressed, triggering the export of HMGB1 from the nucleus and cytoplasm, without undergoing cell death. Non-pyknotic cells that were HMGB1-negative were not abundant in control, undamaged retinas or in retinas at 1hr after NMDA-treatment (Fig. 4a,b,g). By comparison, non-pyknotic cells that were HMGB1-negative were significantly more abundant in INL at 4hrs and were further increased at 24hrs after treatment (Fig. 4c,d,g). Collectively, these data indicate that there is a rapid export of HMGB1 from the nuclei of damaged/stressed neurons in the INL, followed by the accumulation of pyknotic and non-pyknotic cells that are negative for HMGB1. These findings are consistent with the hypothesis that HMGB1 is exported from the nuclei and secreted/released by damaged, stressed or dying cells to provide a “danger” signal to nearby cells.

We next investigated whether transcription factors (TFs) are exported from the nuclei of damaged retinal neurons. Different TFs are known to be exported out the nuclei of stressed or damaged cells (Shukla and Mishra, 2018; Park et al., 2023). Sox2, Islet1 and Pax6 are normally present in the nuclei of cholinergic amacrine cells (Fig. 5a,b,f,g) (Elshatory et al., 2007; Fischer et al., 2007, 2010; Stanke et al., 2008). At 1hr after NMDA-treatment, we detected many Sox2-positive cells in the proximal INL at the IPL border wherein Sox2 was detected in the cytoplasm and nucleus (Fig. 5a). These cells often had diminished levels of Sox2 and HMGB1 in the nucleus, but did not appear to be pyknotic based on DAPI labeling (Fig. 5c-e). These cells were more abundant in peripheral regions compared to central regions of the retina (Fig. 5e). Amacrine cells with Sox2 in the cytoplasm were not observed at 4 or 24 hrs after NMDA-treatment (not shown). We did not observe Sox2 exported out of the nuclei of MG between 1 and 24 hrs after treatment (Fig. 5a-e); however, Sox2 and Sox9 are always observed in the cytoplasm of MGPCs in M-phase when the nuclear envelope is absent (Zelinka et al., 2016; Palazzo et al., 2020; Campbell et al., 2023; Taylor et al., 2024).

**Figure 5:**
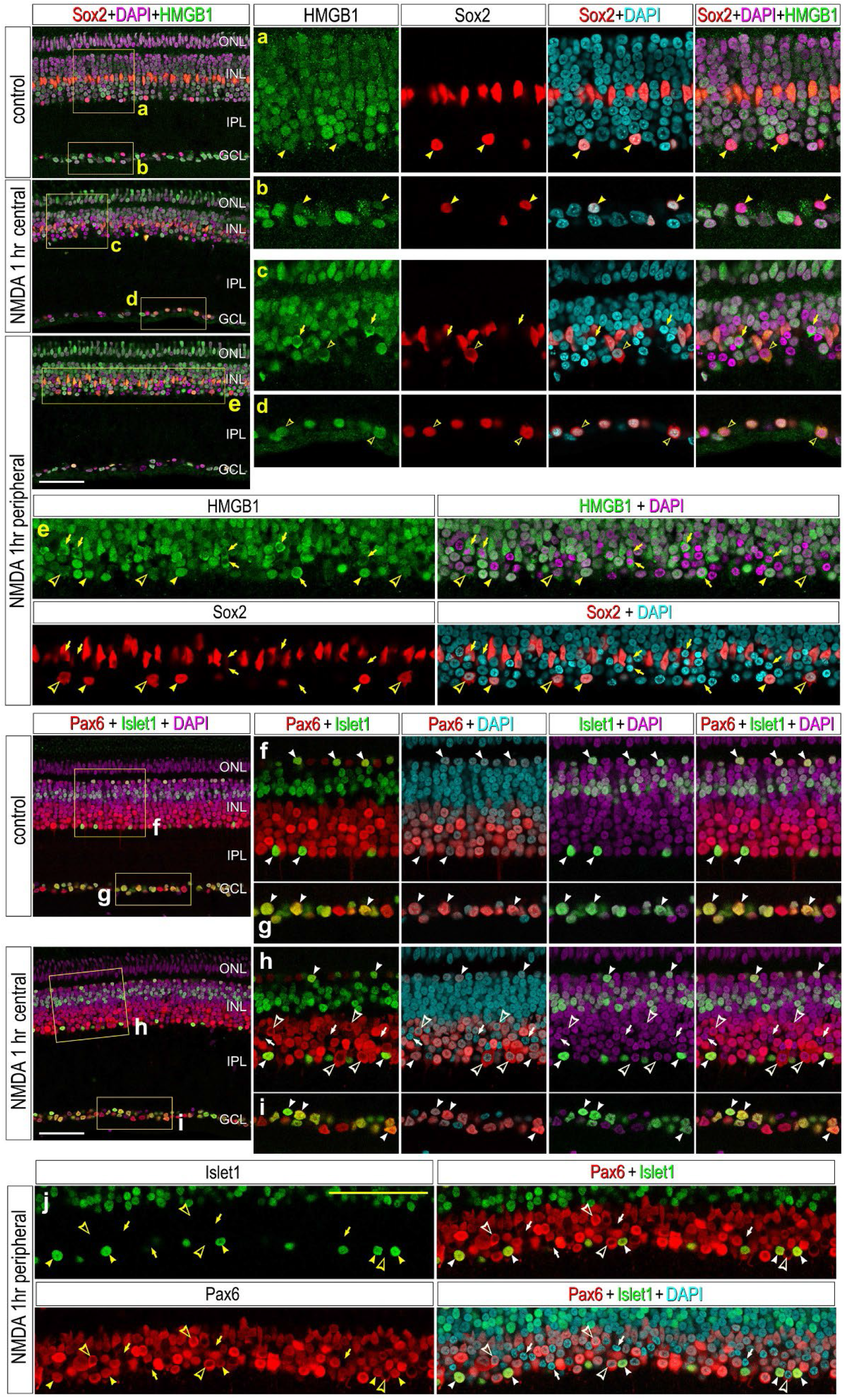
The transcription factors Sox2 and Pax6, but not Islet1, are exported out the nuclei of neurons in damaged retinas. Sections of the retina were obtained from eyes injected with saline (**a,b**) or 1hr after NMDA-treatment (**c-e**). Sections were labeled with DAPI (cyan or magenta) and antibodies to HMGB1 (green; **a-e**), Sox2 (red; **a-e**), Pax6 (red; **f-j**), or Islet1 (green; **f-j**). Confocal microscopy was used to obtain single optical sections that were less than 2 µm thick. Images were obtained from central (**a-d, f-i**) and peripheral (**e,j**) regions of the retina. Solid arrowheads indicate the cholinergic amacrine cells with HMGB1 and Sox2 restricted to the nucleus. Hollow arrowheads indicate cholinergic amacrine cells with Sox2 and HMGB1 in the peri-nuclear cytoplasm. Arrows indicate DAPI-intense nuclei with HMGB1 exported out of the nuclei into the peri-nuclear cytoplasm. Calibration bars in panels **b**, **d** and **f** represent 50 µm. Areas indicated by yellow boxes are enlarged 2-fold in adjacent panels. Abbreviations: ONL – outer nuclear layer, INL – inner nuclear layer, IPL – inner plexiform layer, GCL – ganglion cell layer, ns – not significant. NMDA, N-methyl-D-aspartate.

We next probed for Pax6-immunoreactivity, which is normally present at high levels in the nuclei of amacrine and ganglion cells, and at lower levels in the nuclei of horizontal cells and MG (Fig. 5f,g) (Edqvist et al., 2006; Fischer et al., 2007). In normal retinas, Pax6-immunoreactivity was predominantly localized to the nuclei of amacrine cells and MG in the INL (Fig. 5f,g). At 1 hr after NMDA, numerous amacrine cells had diminished or absent Pax6-immunoreactivity in the nucleus and prominent labeling in the cytoplasm in both central and peripheral regions of the retina (Fig. 5f). The biological significance of exporting Sox2 and Pax6 out of the nuclei of amacrine cells remains unknown.

We next investigated whether TFs in bipolar cells were exported out of nuclei in damaged retinas, because NMDA-treatment is known to damage and destroy a few bipolar cells (Fischer et al., 1998, 2004). Accordingly, we probed for orthodenticle homeobox 2 (Otx2) and Islet1 homeodomain TFs which are known to be expressed by bipolar cells in the chick retina (Trimarchi et al., 2008; Westenskow et al., 2010) (Fig. 5f,g). At 1 hr after NMDA, neither Otx2 or Islet1-immunoreactivities were found outside of the nuclei of bipolar cells (Fig. 5g; Supplemental Fig 3). In addition to expression in bipolar cells, Otx2 is expressed by photoreceptors (Trimarchi et al., 2008; Westenskow et al., 2010) and Islet1 is expressed by cholinergic amacrine cells (Stanke et al., 2008) and about 50% of the horizontal cells (Fischer et al., 2007). At 1 hr after NMDA, Otx2 was not observed outside of the nuclei of photoreceptors (supplemental Fig. 3) and, unlike Sox2, Islet1 and Pax6 were not observed outside of the nuclei of cholinergic amacrine cells (Fig. 5f-j).

### Effects of exogenous HMGB1 and CXCL12 on undamaged retinas

We next investigated whether intravitreal injection of CXCR4 agonist (ATI-2341) or recombinant human HMGB1 or recombinant mouse CXCL12 activated cell signaling in retinal cells. In retinas from eyes injected with vehicle, we observed pS6 (a readout of mTor-signaling) in putative RGCs in the GCL. Levels of pS6 were low or absent in the MG in control retinas (Fig. 6a). A single injection of CXCR4 agonist (ATI-2341) or rCXCL12 induced significant accumulations of pS6 in Sox2+ MG (Fig. 6a,b). A single injection of rHMGB1 had no significant effects of cell signal (Fig. 6b). By comparison, a single intraocular injection of rHMGB1/rCXCL12 resulted a significant accumulation of pS6 in MG with Sox2^+^ nuclei (Fig. 6a,b). These findings are consistent with patterns of expression for CXCR4 in resting MG.

**Figure 6:**
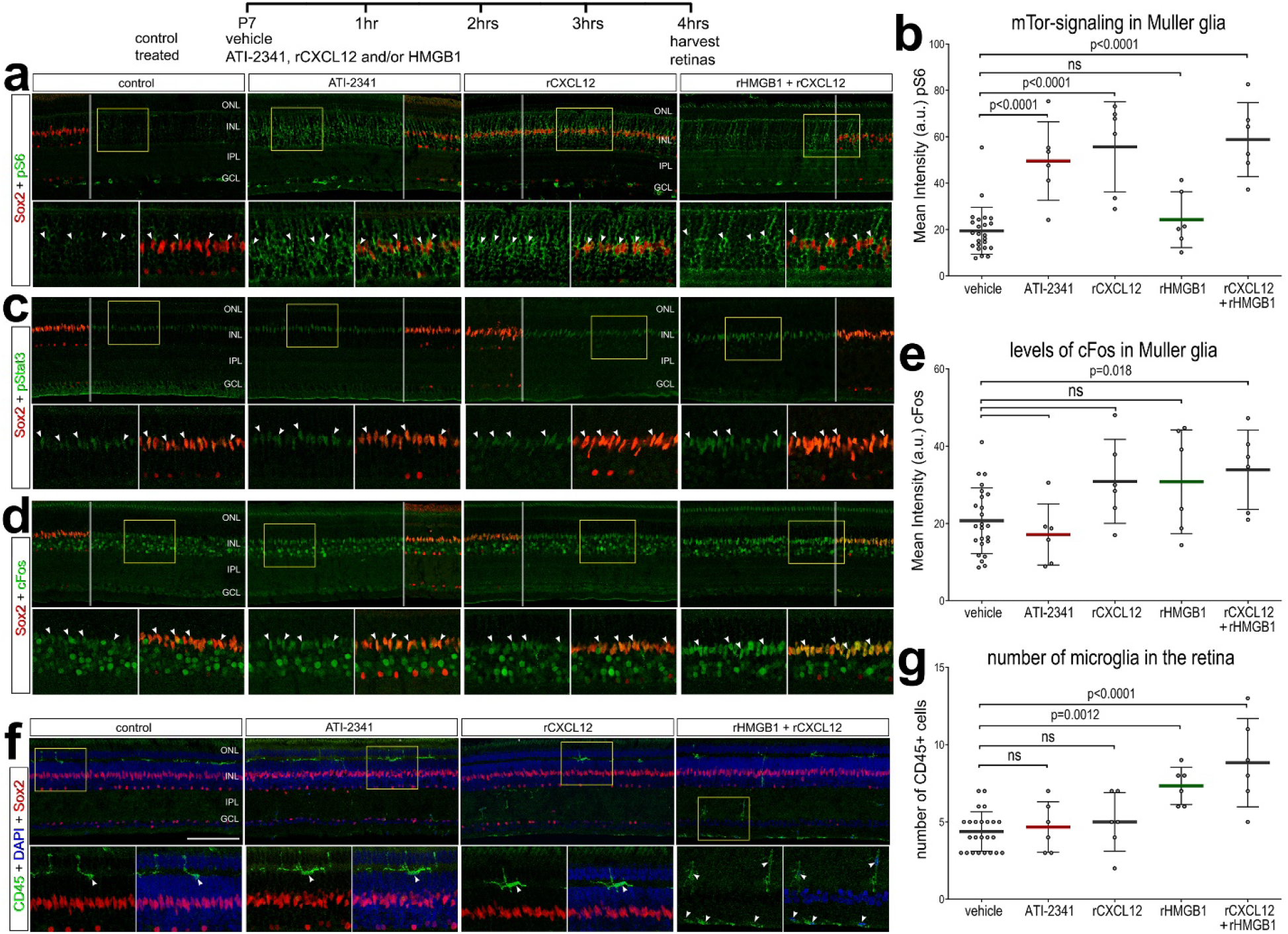
CXCR4 agonist, recombinant HMGB1 and CXCL12 activate mTor-signaling and cFos in MG and the accumulation of microglia. Eyes were injected with vehicle (control) or ATI-2341 (CXCR4-agonist), rCXCL12 alone, rHMGB1 alone or HMGB1+CXCL12 and retinas harvested 4 hrs later. Sections of the retina were labeled with antibodies to Sox2 (red) and pS6 (green) or pStat3 (green) cFos (green) or CD45 (red). Arrow-heads indicate the nuclei of MG. Areas that are boxed-out in yellow are enlarged two-fold in the panels immediately below. Calibration bar in panel **f** represents 50 µm and applies to **a**, **c** and **d**. Areas indicated by yellow boxes are enlarged 2-fold in the panels below. The histograms in **b, e** and **g** illustrate the mean (bar ± SD), each dot represents one biological replicate. Significance of difference was determined by a one-way ANOVA (p<0.0001) and Šidák’s multiple comparisons test (p-values). Abbreviations: ONL – outer nuclear layer, INL – inner nuclear layer, IPL – inner plexiform layer, GCL – ganglion cell layer, ns – not significant. NMDA, N-methyl-D-aspartate.

We next probed for the activation of Jak/Stat signaling (pStat3), which is known to impact the formation of MGPCs in the retinas of fish, chicks and mice (Wan et al., 2014; Todd et al., 2016; Jorstad et al., 2020). In retinas from eyes injected with vehicle, we observed low levels of pStat3 (a readout of Jak/Stat-signaling) in MG with Sox2^+^ nuclei (Fig. 6c). There was no significant accumulation of pStat3 in the nuclei of Sox2+ MG 4 hrs after treatment with ATI-2341, rCXCL12, rHMGB1 or the combination of rCXCL12 and rHMGB1 (Fig. 6c and not shown). We next probed for upregulation of cFos, which is known to activate in MG following different treatments including damage or growth factors such as IGF1 or FGF2 (Fischer et al., 2009a; b). We found that the combination of rCXCl12 and rHMGB1 resulted in a significant increase in cFos expression in MG (Fig. 6d,e). By comparison, ATI-2341, rCXCL12 alone or rHMGB1 alone did not significantly upregulate cFos in MG (Fig. 6d,e).

Consistent with patterns of expression of CXCR4, a single intraocular injection of CXCR4 agonist (ATI-2341) or rCXCL12 had no effect on the abundance of microglia, but the microglia appeared to be more reactive, with elevated levels of CD45) or abundance of microglia (n=6) (Fig. 6f,g). By comparison, a single intraocular injection of rHMGB1 resulted in a small but significant increase in numbers of microglia. Interestingly, the combination of rCXCL12 and rHMGB1 further increased the number of microglia within the retina (Fig. 6f,g). Many of these CD45-positive cells appeared to accumulate at the vitread surface of the retina (Fig. 6f,g). This accumulation of microglia at the vitread surface of the retina is similar to responses of these cells to intravitreal delivery of Interleukin 6 or IGF1 (Zelinka et al., 2012).

Collectively, these findings are consistent with the hypothesis that HMGB1 and CXCL12 activate CXCR4 receptors, expressed by MG, to signal through the mTor pathway and cFos. Further, our data suggest that HMGB1-signaling may induce the accumulation activated microglia in the retina; this could be via CXCR4 signaling in MG or directly through TLR2/4 expressed by microglia.

### Inhibition of HMGB1, CXCR4 or the HMGB1/CXCL12 heterocomplex stimulates the formation of MGPCs

Inflachromene has been shown to directly bind to HMGB1 and suppress post-translational modifications to prevent cytoplasmic localization and release (Lee et al., 2014). Plerixafor is a compound that is clinically approved to specifically inhibit CXCR4. LYS2510924 is a small cyclic peptide which is a potent and selective CXCR4 antagonist currently in phase II clinical studies for cancer. LY2510924 specifically blocks CXCL12 binding to CXCR4 with IC50 value less than 1 nmol/L (Peng et al., 2015). The peptide HBP08 is known to competitively antagonize the formation of the HMGB1/CXCL12 heterocomplex and, thereby, prevent activation of CXCR4 (Sgrignani et al., 2021).

We began testing whether inflachromene inhibited the nuclear export of HMGB1 in stressed neurons in NMDA-treated retinas. Inhibition of HMGB1 secretion with inflachromene significantly reduced the number of HMGB1-cytoplasmic profiles in the inner INL at 4 hrs after NMDA-treatment (Fig. 7a-c). We found that inhibition of HMGB1 secretion, inhibition of the HMGB1/CXCL12 heterocomplex or inhibition of CXCR4 significantly increased numbers of proliferating MGPCs in NMDA-damaged retinas (Fig. 8). We identified proliferating MGPCs by co-labeling for Sox2 and incorporation of EdU (Fig. 8a-e) and co-labeling for neurofilament, Sox9 and phospho-histone H3 (Fig. 8f-j). Two chemically distinct small molecule inhibitors to CXCR4, Plerixafor and LYS2510924, both stimulated the proliferation of MGPCs in damaged retinas (Fig. 8). We found that inflachromene, HPB08, Plerixafor and LYS2510924 had no effects upon numbers of dying cells or the reactivity, accumulation or proliferation of microglia in damaged retinas (Supplemental Fig. 4).

**Figure 7:**
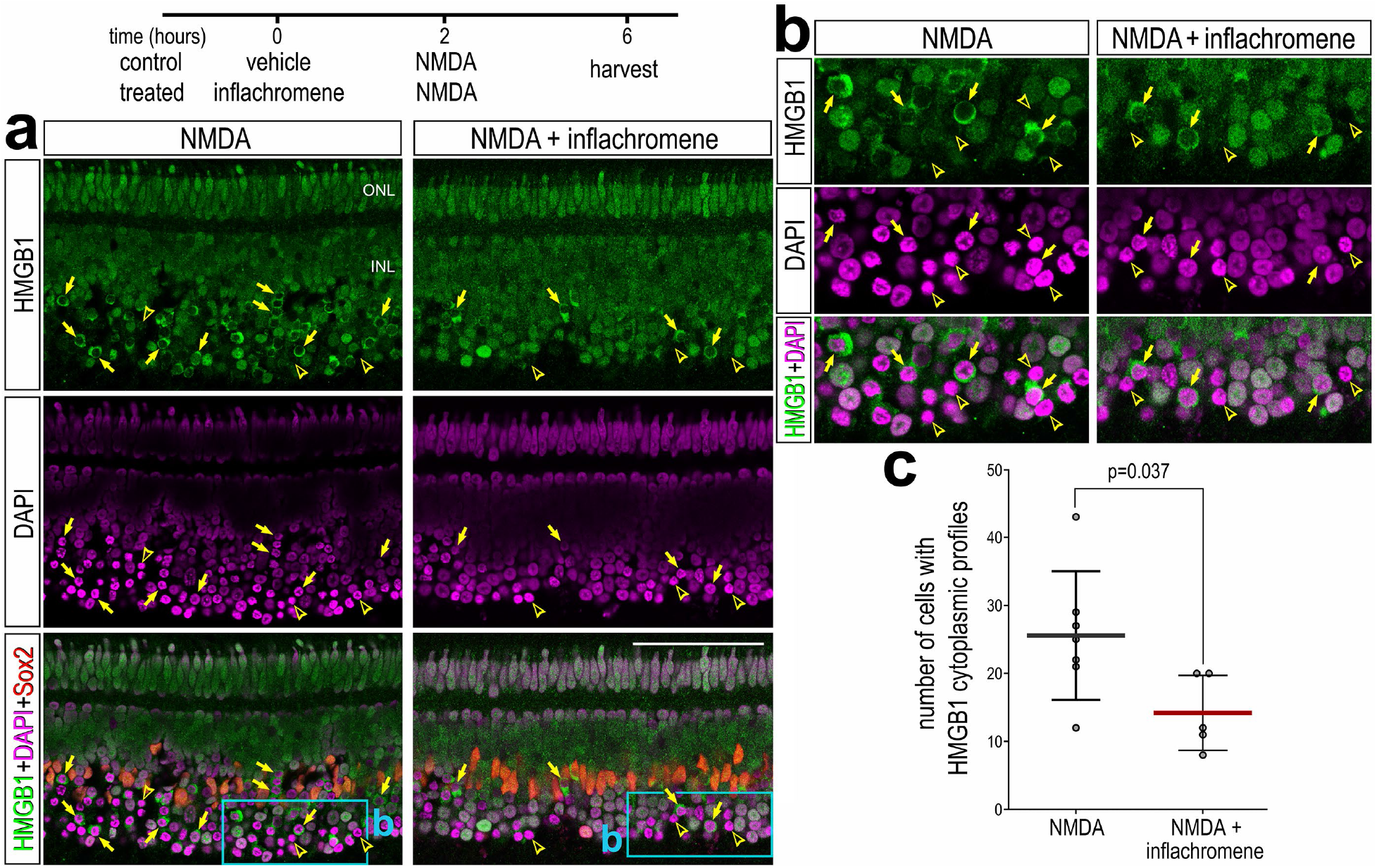
Inflachromene inhibits the nuclear export of HMGB1 in amacrine cells in NMDA-treated retinas. Sections of the retina were obtained from eyes injected with saline or NMDA at 1 hr after treatment (**a,b**). Sections were labeled with DAPI (magenta) and antibodies to HMGB1 (green; **a,b**) and Sox2 (red; **a**). Confocal microscopy as used to obtain single optical sections that were less than 2 µm thick. Hollow arrows (yellow) indicate DAPI-intense nuclei that are negative for HMGB1 and arrows (yellow) indicate DAPI-intense nuclei with HMGB1 exported out of the nuclei into the peri-nuclear cytoplasm. Calibration bar in panels **a** represents 50 µm. Areas indicated by cyan rectangles in **a** are enlarged 2-fold in **b**. The histogram in **c** illustrates the mean (bar ± SD), each dot represents one biological replicate. Significance of difference (p-values) was determined by using an unpaired t-test. Abbreviations: ONL – outer nuclear layer, INL – inner nuclear layer, IPL – inner plexiform layer, GCL – ganglion cell layer, NMDA, N-methyl-D-aspartate.

**Figure 8:**
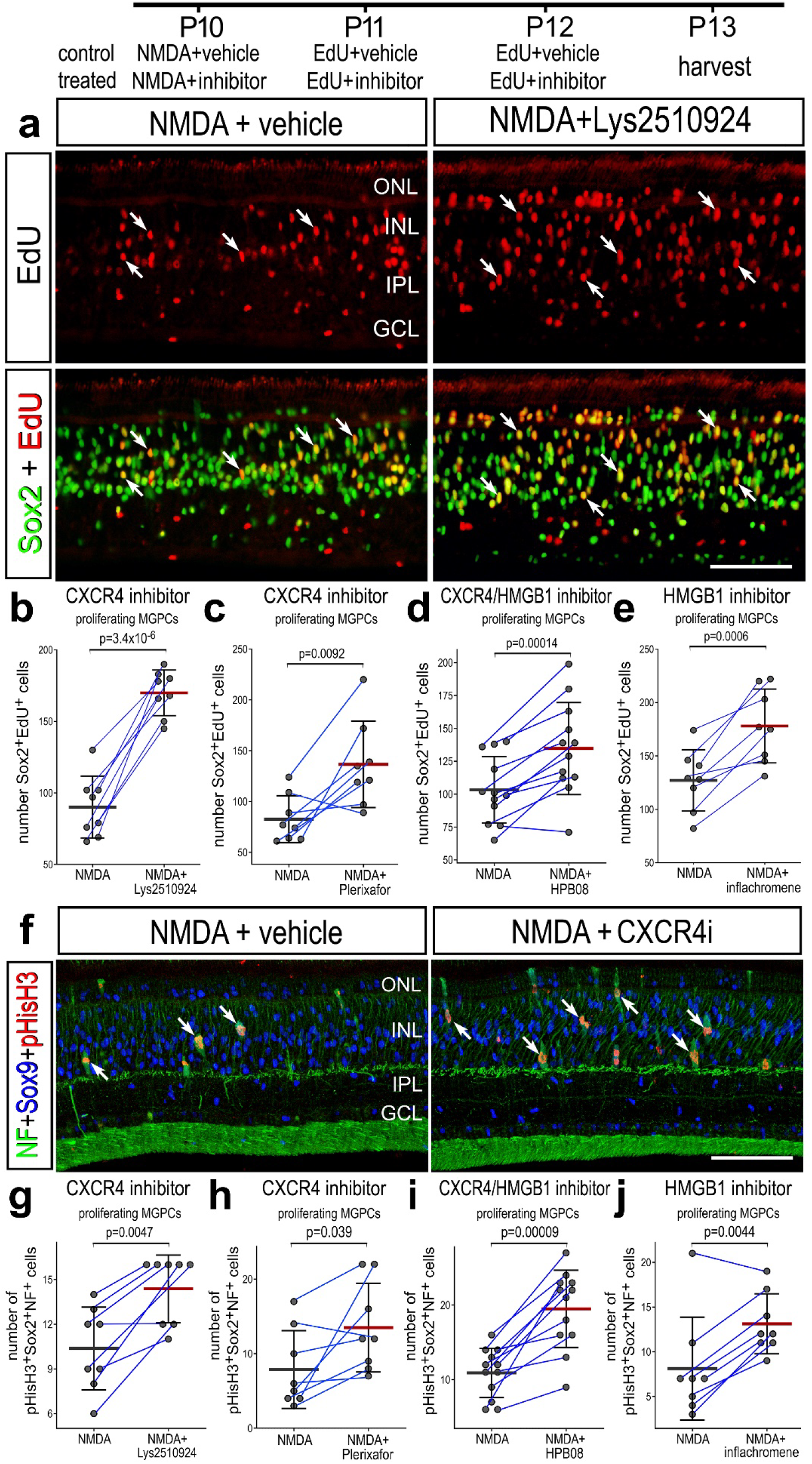
Inhibition of HMGB1 secretion, HMGB1/CXCL12 heterocomplex or CXCR4 stimulates the formation of proliferating MGPCs. Drugs included Lys2519024 (CXCR4 antagonist), Plerixafor (CXCR4 antagonist), inflachromene (inhibitor of HMGB1 secretion), and HPB08 (inhibitor of the HMGB1/CXCL12 heterocomplex). Eyes were injected with NMDA+vehicle (control) or NMDA+inhibitor (treated) at P10, EdU+vehicle or inhibitor at P11 and P12, and retinas harvested at P13, 24 hrs after the last injection. Vertical sections of the retina were labeled for EdU-incorporation and/or antibodies to Sox2, Sox9, neurofilament and phospho-histone H3 (pHisH3). Arrows indicate the nuclei of MGPCs labeled for EdU and Sox2 (**a**) or pHisH3, Sox9 and neurofilament (**f**). The calibration bars in **a** and **f** represent 50 µm. Histograms illustrate mean (bar ± SD) numbers of MGPCs (Sox2+EdU; **b-e** or NF+Sox9+pHisH3; **g-j**). Each dot represents one biological replicate, and blue lines connect replicates from control and treated retinas from one individual (n= 8 to 12). Significance of difference (p-values) was determined by using a paired t-test. Abbreviations: ONL – outer nuclear layer, INL – inner nuclear layer, IPL – inner plexiform layer, GCL – ganglion cell layer.

We next tested whether CXCR4 agonist (ATI-2341) influenced the formation of MGPCs in damaged retinas. We found that CXCR4 agonist had no significant effect upon numbers of proliferating MGPCs (Supplemental Fig. 5). Since ATI-2341 activated mTor and Jak/Stat cell signaling in MG in undamaged retinas, demonstrating an expected biological effect, it is possible that signaling at CXCR4 is saturated in damaged retinas and, therefore, further activation by an agonist had no effect upon MGPC formation. Collectively, these findings indicate that inhibition of HMGB1 secretion, inhibition of the HMGB1/CXCL12 heterocomplex or inhibition of CXCR4 all potently stimulate the formation of proliferating MGPCs in damaged retinas, while not affecting cell death or microglial abundance or reactivity.

### Inhibition of HMGB1 and neuronal differentiation

We next investigated whether inhibition of HMGB1 nuclear export influenced the neurogenic potential of MGPCs. When inflachromene is applied with NMDA and on the 2 days following NMDA, before the majority of MGPCs re-enter the cell cycle (Fischer and Reh, 2001; Fischer, 2005), we found a significant increase in the number of newly generated neurons that were labeled for EdU and HuC/D (Fig. 9a-d). These cells were commonly found in the INL and were occasionally found in the GCL (Fig. 9b,c).

**Figure 9:**
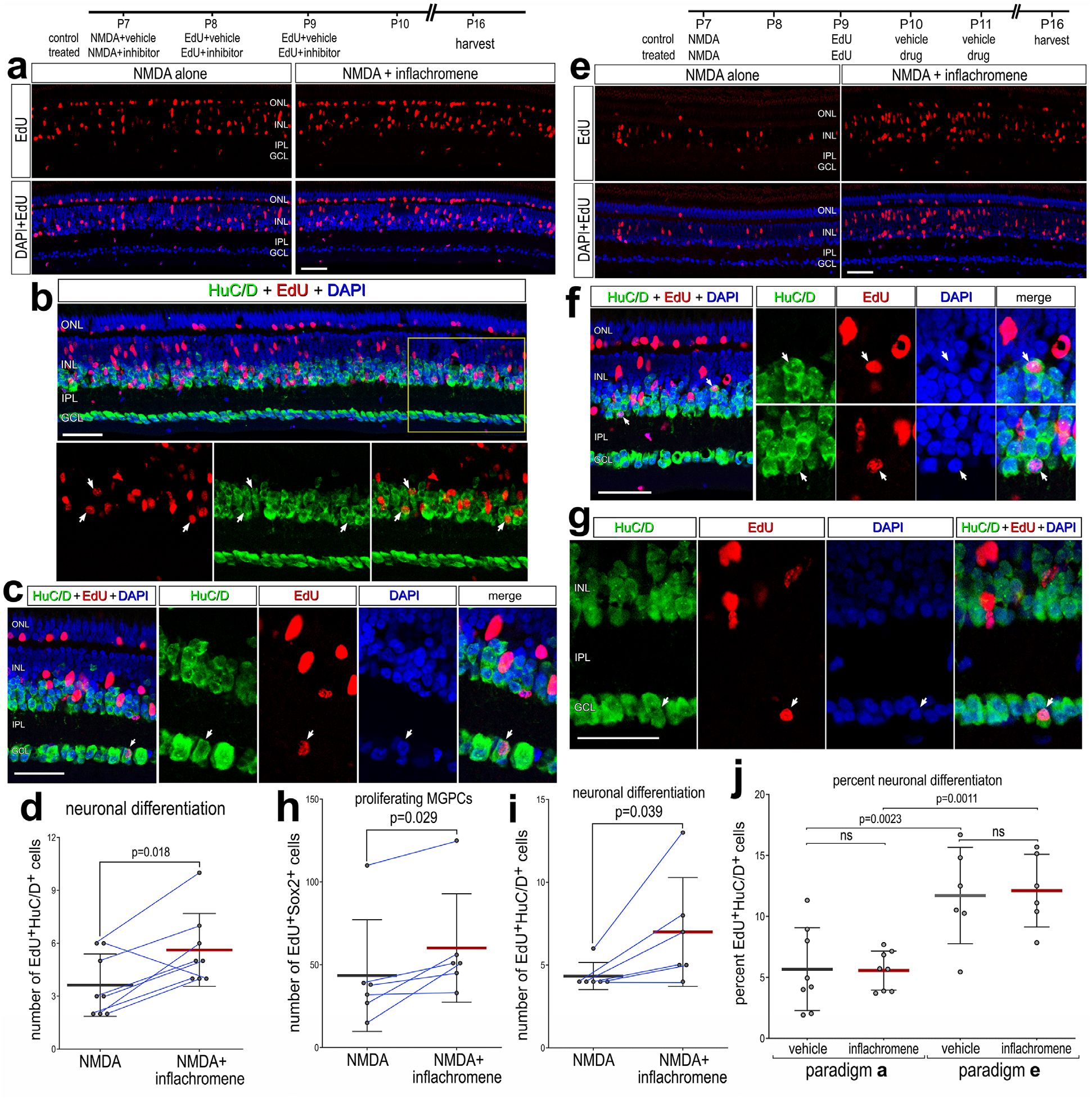
Inhibition of HMGB1 secretion enhances neuronal differentiation from MGPCs. Eyes were injected with NMDA at P7, EdU plus vehicle or drug at P8 and P9, and retinas harvested at P16 (**a-d**). Alternatively, eyes were injected with NMDA at P7, EdU at P9, vehicle or inflachromene at P10 and P11, and retinas harvested at P16 (**e-j**). Retinas sections were labeled for EdU (red; **a-c, e-g**), DAPI (blue; **a-c**, **e-g**) and antibodies to HuC/D (green; **b,c,f,g**). Arrows indicate the nuclei of regenerated neurons that are labeled for EdU and HuC/D. Calibration bars in panels **a-c** and **e-g** represent 50 µm. Histograms represent the mean (bar ± SD) and each dot represents one biological replicate (blue lines connect data points from control and treated retinas from the same individual) (**d,h,i,j**). Significance of difference (p-values) was determined by using a paired t-test (**d,h,i**) or one-way ANOVA (p=0.0002) with post-hoc Šidák’s test for multiple comparisons (**j**). Abbreviations: ONL – outer nuclear layer, INL – inner nuclear layer, IPL – inner plexiform layer, GCL – ganglion cell layer, ns – not significant, NMDA, N-methyl-D-aspartate, EdU-5-Ethynyl-2-deoxyuridine.

When EdU was applied at 2 days and inflachromene was applied at 3 and 4 days after NMDA, we found a significant increase in the number of proliferating MGPCs (Fig. 9e,h). In addition, we found a significant increase in the abundance of newly generated neurons (Fig. 9f,g,i). These cells were commonly found in the INL and include some found in the GCL (Fig. 9g). Treatment with inflachromene did not significantly affect the percentage of EdU-HuC/D neurons generated by MGPCs regardless of when the EdU and inflachromene were applied after NMDA (Fig. 9j). However, we found a significant increase in the percentage of EdU-HuC/D neurons when the inflachromene was applied starting at 3 days after NMDA compared to drug-treatment starting with the NMDA (Fig. 9h). Collectively, these findings indicate that the increase in the abundance of new neurons in damaged retinas treated with inflachromene was secondary to increases in the abundance of proliferating MGPCs. In addition, these findings suggest that the MGPCs that proliferate at later times after NMDA-treatment have increased neurogenic potential.

### Inhibition of HMGB1 secretion and CXCR4 had no effect on MGPCs formation in undamaged retinas

If HGMB1 is released from damaged neurons to form a complex with CXCL12 to activate CXCR4, then inhibition of HMGB1 secretion or inhibition of CXCR4 should have no effect on the formation of MGPCs in undamaged retinas. MGPCs are known to form in undamaged retinas that are treated with 3 consecutive daily injections of insulin and FGF2 (Fischer et al., 2002, 2014). We began by analyzing scRNA-seq libraries or chick retinas treated with 2 or 3 consecutive daily doses of insulin and FGF2; these libraries were generated and analyzed previously (Hoang et al., 2020; Palazzo et al., 2020; Campbell et al., 2021b). UMAP embedding of retina cells revealed discrete clusters of retinal neurons that were embedded together based on major cell class regardless of treatment, whereas MG formed distinct clusters that were segregated based on treatment with 2 or 3 doses of insulin and FGF2 (Fig. 10a-c). Resting MG, activated MG and MGPCs were identified based on expression of *GLUL, RLBP1, PMP2, S100A6, CDK1* and *TOP2A* (Fig. 10d-f). Patterns of expression of *HMGB1, HMGB2, CXCL12* and *CXCR4* were consistently seen in aggregate libraries of damaged chick retinas. *HMGB1* and *HMGB2* were highly expressed by retinal neurons and MGPCs (Fig. 10g,h,k). *CXCL12* was expressed by bipolar and amacrine cells (Fig. 10i). *CXCR4* was highly expressed by resting MG and was significantly downregulated by MG treated with 2 or 3 doses of insulin and FGF2 (Fig. 10j,k).

**Figure 10:**
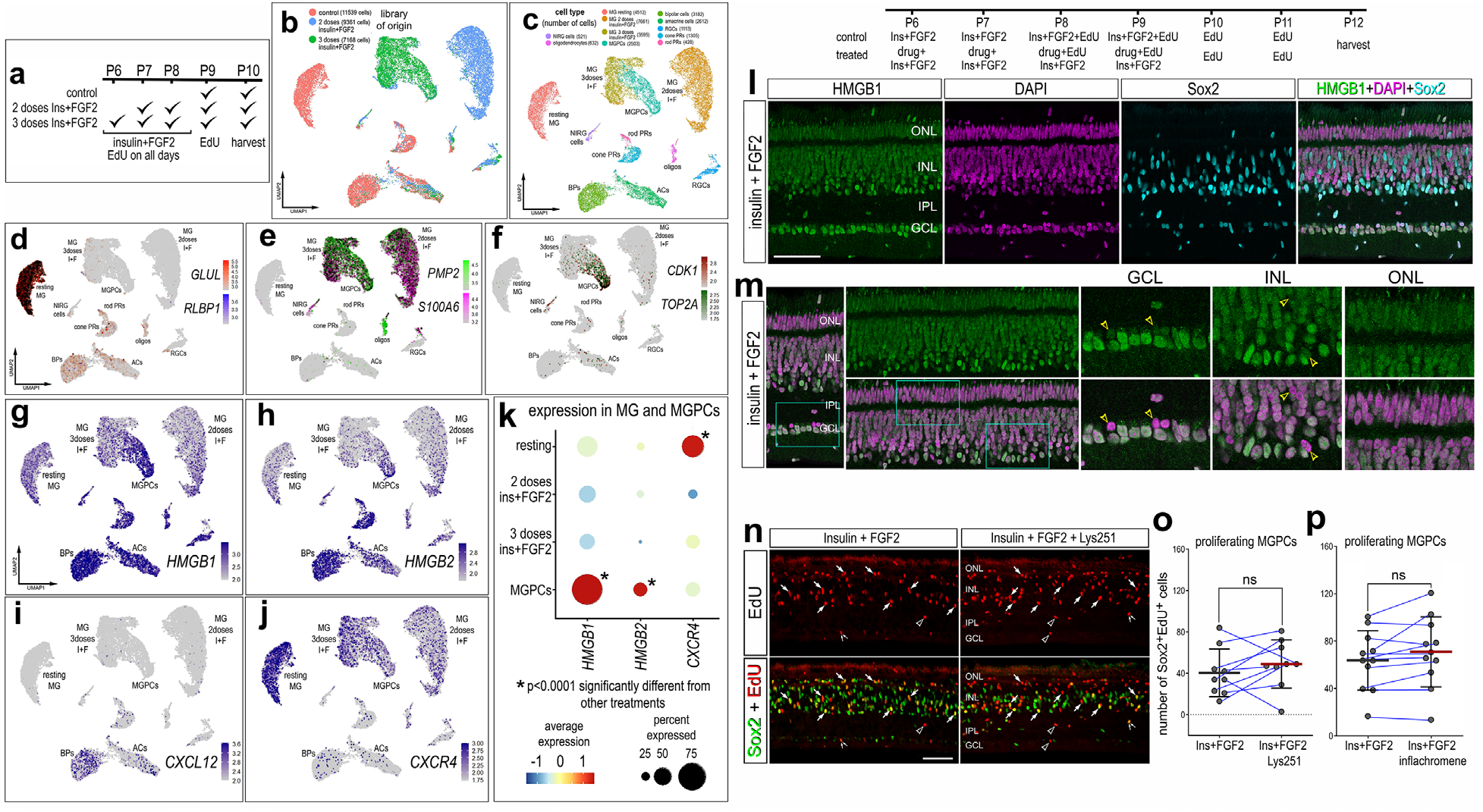
Insulin and FGF2 treatment downregulates *CXCR4* in MG, HMGB1 remains confined to nuceli, and the formation of MGPCs is not affected by inflachromene or CXCR4 inhibitor. scRNA-libraries were generated and characterized previously (Palazzo et al., 2020; Campbell et al., 2021). scRNA-seq libraries were generated from cells obtained from undamaged retinas and from retinas treated with 2 or 3 consecutive daily injections of insulin and FGF2 (**a**). Clusters of MG and MGPCs were identified based expression of resting markers (*GLUL* and *RLBP1*; **d**), activated markers (*PMP2* and *S100A6*; **e**) and proliferation (*CDK1* and *TOP2A*; **f**). Feature plots in **g-j** illustrate patterns and levels of expression of *HMGB1, HMGB2, CXCL12,* and *CXCR4* in UMAP clustered retinal cells. Dotplots in **k** illustrate the average expression (heatmap) and percentage of expressing cells (dot size) for *HMGB1, HMGB2,* and *CXCR4* in MG and MGPCs. Wilcoxon Rank Sum Test with Bonferroni correction was used to determine the significance of difference (*p<<0.0001; supplemental table 1). Eyes were injected with insulin+FGF2 plus vehicle or drug at P6 and P7, insulin+FGF2+EdU plus vehicle or drug P8 and P9, EdU at P10 and P11, and retinas harvested at P12 (**l-p**). Retinal sections were label with DAPI (magenta; **I,m**) and antibodies to HMGB1 (green; **l,m**), Sox2 (cyan; **l**; green **n**), or EdU (red; **n**). Yellow hollow arrow-heads indicate DAPI-positive nuclei with little or no HMGB1-immunoreactivity (**m**). Cyan-colored boxes in **m** are enlarged 2-fold in the panels to the right. Arrows indicate the nuclei of MGPCs labeled for Sox2 and EdU (**n**). Histograms (**o,p**) illustrate the mean (bar ± SD), each dot represents one biological replicate, and blue lines connect counts from control and treated retinas from one individual. Significance of difference (p-values) was determined by using a paired t-test. Abbreviations: ONL – outer nuclear layer, INL – inner nuclear layer, IPL – inner plexiform layer, GCL – ganglion cell layer, ns – not significant, NMDA, N-methyl-D-aspartate, EdU-5-Ethynyl-2-deoxyuridine.

We did not find any cells with HMGB1 exported out of the nucleus and into the cytoplasm in retinas treated with insulin and FGF2 (Fig. 10l,m). This is consistent with the fact that neurons are not damaged or stressed in response to insulin and FGF2. Accordingly, we found that inhibition of HMGB1 trafficking had no effect on the formation of MGPCs (Fig. 10n-p). Similarly, inhibition of CXCR4 had no effect on numbers of proliferating MGPCs in retinas treated with insulin+FGF2 (Fig. 10n-p). The failure of the CXCR4 inhibitor to influence MGPC proliferation may occur because *CXCR4* is downregulated by insulin+FGF2 or because HMGB1 is not exported out of nuclei in undamaged neurons. Alternatively, it is possible that MGPCs are “permitted” to proliferate in response to insulin+FGF2 because *CXCR4* is downregulated to prevent CXCR4 signaling, which suppresses the reprogramming of MG.

### Microglia influence the expression of HMGB’s and CXCR4 in MG

Reactive microglia are known to impact the formation of proliferating MGPCs in the zebrafish retina (Huang et al., 2012; White et al., 2017) and suppress Ascl1-mediated reprogramming of MG into neuronal cells in the rodent retina (Todd et al., 2020; Palazzo et al., 2022). In the chick retina, the ablation of activated microglia blocks the formation of MGPCs in damaged and undamaged retinas (Fischer et al., 2014). However, the failed formation of MGPCs in damaged retinas missing microglia can be rescued by activation of NFkB, exogenous TNF (Palazzo et al., 2020) or exogenous FGF2 or HBEGF (El-Hodiri et al., 2023). Accordingly, we tested whether the expression of HMGB’s or CXCR4 was influenced by microglia and whether inhibition of HMGB1/CXCL12-CXCR4-signaling could rescue the formation of MGPCs in damaged retinas missing microglia.

Microglia were ablated by applying clodronate liposome. A single intravitreal injection of clodronate liposomes destroys >95% of microglia within 3 days of treatment, whereas numbers of MG are unaffected (Zelinka et al., 2012; Fischer et al., 2014). We analyzed scRNA-seq libraries of normal and NMDA-damaged chick retinas with and without microglia, as described previously (Palazzo et al., 2020; El-Hodiri et al., 2023). The aggregate dataset (41,162 total cells) included scRNA-seq libraries from (i) control undamaged retina, (ii) undamaged retinas with microglia depleted, (iii) NMDA-damaged retinas at 24 hrs after treatment, and (iv) NMDA-damaged retinas at 24 hrs after treatment with microglia depleted (Fig. 11a-c). UMAP plots revealed distinct clusters of cell types, with neuronal types clustered together regardless of treatment and MG formed distinct clusters that correlated with treatments (Fig. 11a,b). Clusters of retinal cells were identified based on cell-distinguishing markers, as described in Materials and Methods. Consistent with patterns of expression seen in other scRNA-seq libraries, *HMGB1, HMGB2* and *HMGB3* were widely expressed by retinal neurons and MG, with the exception of little or no expression of *HMGB2* and *HMGB3* in RGCs (Fig. 11d,e). *CXCR4* was predominantly expressed by MG (Fig. 11e). The depletion of microglia caused MG to significantly downregulate *CXCR4* in damaged and undamaged retinas (Fig. 11f). By comparison, levels of *HMGB1* were significantly higher in MG in damaged retinas when the microglia were ablated (Fig. 11f). Inhibition of HMGB1 secretion or inhibition of CXCR4 had no effect on numbers of dying cells or proliferating MGPCs labeled with EdU or antibodies to phospho-histone H3 in NMDA-damaged retinas in the absence of microglia (Fig. 11g-i). These inhibitors likely had no effect upon the formation of proliferating MGPCs because *CXCR4* was significantly downregulated by MG in normal and damaged retinas when the microglia were ablated. It is also possible that HMGB1 inhibitor had no effect on MGPC proliferation because TLR2/4-receptor-expressing cells, the microglia, were absent.

**Figure 11:**
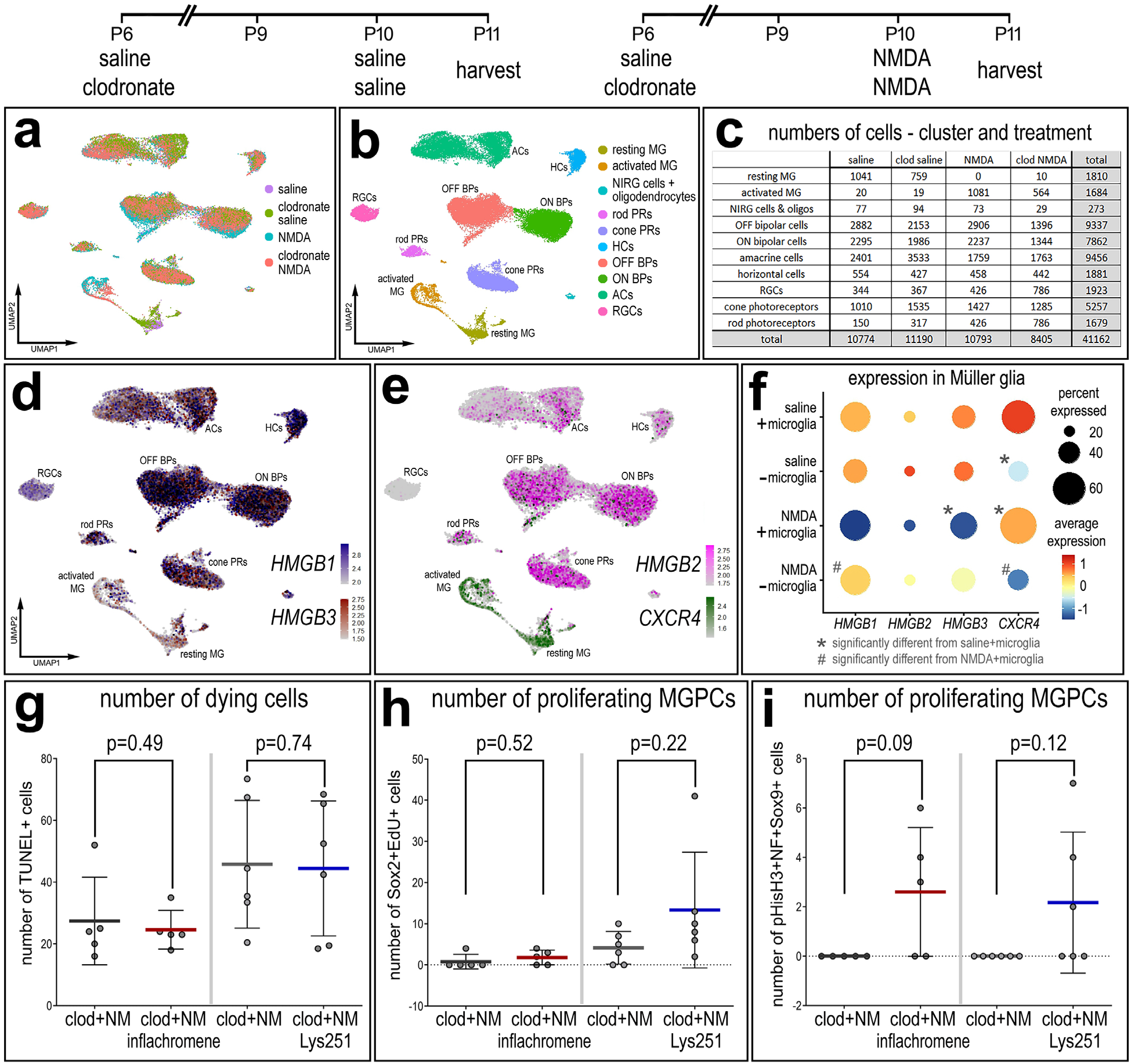
Microglia are required for inhibition of HMGB1/CXCR4 signaling to stimulate MGPC proliferation. scRNA-libraries were generated and analyzed previously (El-Hodiri et al., 2023). Histograms (**b-e**, **g-j**) illustrate the mean (bar ± SD), each dot represents one biological replicate, and blue lines connect counts from control and treated retinas from one individual. Significance of difference (p-values) was determined by using a paired t-test. Feature plots in **c** illustrate patterns and levels of expression of *HMGB1, HMGB2, CXCL12, TLR2* and *TLR4* in UMAP clustered retinal cells. The violin plot in **d** illustrates levels and percent expression in MG in control retinas and at different times after NMDA-treatment. Dotplots in **e** illustrate the average expression (heatmap) and percentage of expressing cells (dot size) for *HMGB1, HMGB2, TLR2, TLR4* and/or *CXCL12* in whole retina, microglia, INL neurons or RGCs.

## Discussion

We find dynamic patterns of expression of CXCR4 and HMGB’s in different types of retinal cells following retinal damage. Although static levels of mRNA are not predictive of protein levels and function, dynamic changes of mRNA levels are strongly correlated with changes in protein levels and function (Liu et al., 2016). These dynamic changes in HMGB’s, CXCL12 and associated receptors in distinct types of retinal cells, suggest that these cells are impacted by signaling through CXCR4 and TLR2/4. Since HMGB1 is known to be released/secreted by damaged or stressed cells to elicit inflammatory responses (Chen et al., 2022), it is expected that HMGB1 signaling through CXCR4 and TLR2/4 elicits pro-inflammatory responses in damaged retinas.

We propose that HMGB1-mediated pro-inflammatory signaling in damaged retinas is manifested by HMGB1 release from damage/stressed neurons and complexing with CXCL12 to activate CXCR4 in MG, whereas activation of TLR4 in microglia may be minimal. Consistent with this hypothesis, we found that exogenous rHMBG1 and rCXCL12 selectively activated cell signaling in MG. We found that inhibition of HMGB1 release/secretion and inhibition of the HMGB1/CXCL12 heterocomplex had no effects upon the reactivity, accumulation or proliferation of microglia in damaged retinas. CXCR4 agonist had no effects upon microglia in undamaged retinas, consistent with patterns of expression from scRNA-seq datasets indicating the MG in the chick retina are the primary cell type that expresses *CXCR4*. Inhibition of HMGB1 release had no significant effects upon the accumulation of microglia in damaged retina. In undamaged retinas, by comparison, rHMGB1± rCXCL12 resulted in a significant accumulation of microglia. It is possible that HMGB1 acts at TLR2/4 to directly impact microglia and secondarily influence MG. In the mouse retina, intravitreal delivery of Tlr2 agonist resulted in the activation of the NFkB-pathway in MG in the mouse retina, but only when microglia were present in the retina (Palazzo et al., 2023). In the chick retina, however, *TLR2* is not highly expressed by microglia and *TLR4* appears to be downregulated by microglia in damaged retinas (Fig. 3). Collectively, these findings suggest that HMGB1-mediated activation TLR2/4 receptors on microglia may not occur, or is superseded by different signals, in damaged chick retinas. However, our findings indicate that microglia provide signals to MG to influence levels of expression of *CXCR4*. The identity of these microglia-derived signals remains unknown. Further studies are required to investigate whether activation of TLR2/4 on microglia is involved in the responses of glia to neuronal damage and the formation of MGPCs.

Pro-inflammatory signaling acts to influence the regenerative capacity of MG after damage in different vertebrate species (Bhattacharya et al., 2025; Taylor et al., 2025b). Thus, it is important to understand how pro-inflammatory signaling fits into the complex network of pathways that control MG reprogramming. There is evidence to suggest that rapid activation of pro-inflammatory signaling from microglia (namely Il1a, IL1b and TNF) activates NFkB signaling in MG (Todd et al., 2019; Palazzo et al., 2022, 2023). Sustained activation of pro-inflammatory signaling suppressed the proliferation and neurogenic potential of MGPCs (reviewed by (Taylor et al., 2025b))

It is possible that HMGB2 and HMGB3 are released by damaged retinal neurons to drive inflammation. Similar to HMGB1, when released into the extracellular space, HMGB2 and HMGB3 act as Damage-Associated Molecular Patterns (DAMPs) or alarmins, acting as danger signals that trigger immune responses, inflammation, and cellular stress reactions (Yang et al., 2010; Silvin et al., 2022; Jiang et al., 2024). HMGB2 can be released by damaged vascular smooth muscle cells during tissue injury (Zheng et al., 2023). Elevated levels of extracellular HMGB2 have been linked to pulmonary hypertension, femoral artery injury, and abdominal aortic aneurysm by promoting inflammation and ferroptosis (Wu et al., 2023). HMGB2 functions as a DAMP in gliomas, contributing to immune response pathways after radiation therapy (Jiang et al., 2024). While predominantly nuclear, HMGB3 can be released under conditions of severe cellular stress or injury (Chikhirzhina et al., 2024). Extracellular HMGB3 acts as a DAMP, which can activate the NLRP3 inflammasome and trigger inflammatory responses (Chikhirzhina et al., 2024). In glioma cells, HMGB3 has been shown to be released via exosomes (Zhang et al., 2026). HMGB3 is involved in therapy resistance, particularly in ovarian and breast cancer, where it modulates the tumor microenvironment (Zhang et al., 2026). DAMP-related genes including *AGER* not expressed in normal or damaged chick retinas (not shown), thus mechanisms of action remain unknown

There are several possibilities that may underlie the failure of CXCR4 inhibitor and inflachromene to influence the formation of MGPCs in undamaged retinas treated with insulin and FGF2. The failure of CXCR4 inhibitor to influence MGPC proliferation may occur because *CXCR4* is downregulated by treatment with insulin and FGF2. The failure of inflachromene to influence MGPC proliferation may occur because HMGB1 is not exported out of nuclei nor released from undamaged neurons. Another interpretation is that MGPCs are “permitted” to proliferate in response to insulin and FGF2 because *CXCR4* is downregulated to prevent HMGB1/CXCL12-CXCR4 signaling in MG.

### Specificity of drugs and Cell Signaling downstream of CXCR4 in MG

Although the primary amino acid sequences of most cytokines, chemokines and their receptors are not well conserved across vertebrate classes, CXCL12, CXCR4 and HMGB1 are highly conserved. Compared to human CXCL12, chick CXCL12 has 98% positives (identical amino acids plus homologous substitutions) with no gaps. Compared to human CXCR4, chick CXCR4 has 91% positives with no gaps. Similarly, HMGB1 is >95% identical between human and chick proteins. Thus, it is expected that recombinant factors and drugs with well-characterized specificity at human proteins will retain specificity in the chick proteins.

The drugs applied in the current study are expected to inhibit different steps in the HMGB1-mediated inflammatory response, from HMGB1 trafficking out of the nucleus, to formation of the HMGB1/CXCL12 heterocomplex and activation of the CXCR4 receptor (Fig. 1). All of these different inhibitors had consistent effects upon the proliferation of MGPCs, cell death, microglial reactivity and accumulation. If these drugs lacked specificity it would be expected that the antagonists might elicit opposite or inconsistent effects. Further evidence of specificity was demonstrated by inflachromene reduced numbers of amacrine cells with HMGB1 filling the peri-nuclear cytoplasm. These findings are consistent with the notion that inflachromene inhibits the post-translational modifications to export HMGB1 out of the nucleus and out of the cell (Lee et al., 2014).

In the chick retina, CXCR4 is predominantly expressed by MG. Thus, it is expected that an intravitreal delivery of CXCR4 agonist and rCXCL12/rHMGB1 would activate cell signaling in MG. Indeed, we found that CXCR4 agonist and rCXCL12/rHMGB1 activated mTor-signaling in MG. It is likely that different cell signaling pathways, in addition to mTor, are activated downstream of CXCR4. The mTor pathway is known to be required for the formation of MGPCs; mTor-inhibitors potently suppress the formation of MGPCs in the chick retina (Zelinka et al., 2016). Further studies are required to identify the pathways and networks of genes that are downstream of HMGB1/CXCL12-CXCR4 signaling that act to suppress the formation of MGPCs.

The increases in numbers of proliferating MGPCs resulting from inhibition of CXCL12/HMGB1-CXCR4 signaling are much greater than those seen for inhibiting NFkB-(Palazzo et al., 2020), TGFB/Smad2/3-(Todd et al., 2017), glucocorticoid receptors (Gallina et al., 2014) or activating hedgehog-(Todd and Fischer, 2015), BMP/Smad1/5/9 (Todd et al., 2017), Wnt (Gallina et al., 2015a) or Jak/Stat-signaling (Todd et al., 2016). By comparison, inhibition of cell signaling through the mTor pathway potently blocks the proliferation of MGPCs, without affecting MAPK signaling or activation of immediate early genes and overrides the pro-proliferative effects of activating hedgehog-, glucocorticoid-or Wnt-signaling (Zelinka et al., 2016). Intravitreal injection of CXCR4 agonist, rCXCL12 and the combination of rCXCL12 and rHMGB1 resulted in the activation of mTor-signaling in MG, whereas rHMGB1 alone or the combination of rHMGB1+ rCXCL12 stimulated MG to express cFos. We have previously shown that cell signaling through the mTor pathways are required to drive the formation of proliferating MGPCs in the chick retina (Zelinka et al., 2016). Further, upregulation of cFos in MG is consistently associated with treatments that drive the formation of MGPCs (Fischer et al., 2009b; a; Todd and Fischer, 2015; Zelinka et al., 2016). However, in undamaged retinas rCXCL12 and/or rHMGB1 stimulated MG to upregulate pS6 and cFos. It remains unknown how CXCL12 and HMGB1 influence pS6 and cFos in acutely damaged retinas.

### HMGB1 and CXCR4 and neuronal differentiation

Many different signals can influence the neurogenic capacity of MGPCs in the chick retina. Similar to fine-tuning neurogenesis during development, these factors and pathways include Notch-, retinoic acid-, glucocorticoid-, Jak/Stat-signaling and ID factors (reviewed by Taylor et al., 2025). Although we found increased total numbers of newly generated neurons in damaged retinas treated with inflachromene, this treatment also increased total numbers of proliferating MGPCs. When adjusted for total numbers of the proliferating MGPCs, the percentage of differentiating neurons was not significantly affected by inflachromene. In prior studies, we discriminated between effects on proliferation versus differentiation by applying drugs starting at 3 days after NMDA, since MGPCs re-enter the cell cycle at 2 days after NMDA (Fischer and Reh, 2001; Fischer et al., 2004). We have not identified factors or pathways that stimulate the proliferation of MGPCs beyond 3 days after NMDA-treatment (reviewed Taylor et al., 2025). However, current findings suggest that inflachromene can further stimulate the proliferation of MGPCs beyond the 3 day-timepoint.

### Nuclear export of transcription factors in stressed neurons

We found that different TFs are exported out of nuclei of stressed or damaged amacrine cells. Some TFs are transported to the cytoplasm from the nucleus after they regulate gene expression. For example, OTX2 is exported from nuclei and trafficked to lysosomes for degradation (Shukla and Mishra, 2018). Similarly, dynamic changes in the sub-cellular localization of Pax6 regulate activity. For example, in corneal epithelial cells, oxidative stress mediated changes in the expression of Pax6 are associated with changes in sub-cellular localization (Shukla and Mishra, 2018). It should be noted that these studies were conducted in cell culture preparations, which is a stressful condition and may influence trafficking of TFs. Nuclear export of Prospero may be required for degradation and cessation of transcriptional activity (Demidenko et al., 2001). Similarly, export of Pdx1 out of the nucleus and into the cytoplasm can be a fast-acting mechanism to turn-off transcriptional activity once a developmental signal has passed (Wescott et al., 2009). Many homeodomain proteins continuously shuttle between the nucleus and cytoplasm via the CRM1/Exportin pathway; preferential localization to the cytoplasm often targets these proteins for degradation to inactivate transcription (Knauer et al., 2005). We propose that nuclear export of Pax6 or Sox2 in amacrine cells in damaged retinas may occur to turn-off transcriptional activity and may impact survival. Further studies are required to identify the biological significance of rapid nuclear export of Pax6 or Sox2 in amacrine cells.

Some cholinergic amacrine cells selectively export Sox2, but not Pax6 or Islet1 in damaged retinas. In the chick retina, there are 4 types of cholinergic amacrine cells. Type I cells have cell bodies at the INL/IPL border and express Islet1 and Sox2 (Fischer et al., 1998; Stanke et al., 2008). Type II cells have cell bodies displaced to the GCL and express Islet1 and Sox2 (Stanke et al., 2008). Type IIIa cells have somata located toward the middle of the INL and express neurotensin, somatostatin and enkephalin, and the Type IIIb cells have somata toward the middle of the INL, but are negative for neurotensin, somatostatin and enkephalin (Fischer et al., 1998). NMDA-treatment is known to destroy the majority of Type II and IIIa/b cells while sparing the majority of Type I cholinergic amacrine cells (Fischer et al., 1998). We found that Type II cholinergic amacrine cells selectively export Sox2, and this nuclear export does not include Pax6 or Islet1. These findings suggest cell type-specific responses involving nuclear export of different TFs in acutely damaged retinas.

## Conclusions

In summary, our findings indicate that the initiation of glial reactivity occurs, at least in part, via HMGB1 release from stressed neurons in acutely damaged retinas. HMGB1 is rapidly trafficked from the nucleus to the cytoplasm of stressed amacrine cells in damaged retinas, and this trafficking can be suppressed by inflachromene. Secreted HMGB1 likely acts at CXCR4 on MG and possibly TLR4 on microglia to initiate inflammation. Inhibition of HMGB1 release, inhibition of the HMGB1/CXCL12 heterocomplex, and inhibition of CXCR4 potently stimulates the formation of proliferating MGPCs. Future studies are required to investigate whether HMGB1 signaling influences inflammation and MGPCs in the mammalian and fish retinas.

## Data availability

CellRanger output files for Gene-Cell matrices for scRNA-seq data for libraries from saline and NMDA-treated retinas are available through Sharepoint links for embryonic chick retina databases and post-hatch normal and treated chick retina databases: https://osumc.sharepoint.com/sites/Links/FischerLab/Forms/AllItems.aspx scRNA-seq datasets are deposited in GEO (GSE135406, GSE242796) and Gene-Cell matrices for scRNA-seq data for libraries chick retinas treated with saline or NMDA retinas are available through NCBI (GSM7770646, GSM7770647, GSM7770648, GSM7770649). Cell Ranger matrix, features and barcodes files for the snRNA-seq dataset of zebrafish retinas were downloaded through GEO (GSE239410).

## Author Contributions

LEK, OBT, SL, EP, ECH, JW, MG and AT executed experiments, gathered data, and editted the manuscript. HME designed experiments, analyzed data and wrote the manuscript. AJF designed experiments, analyzed data, constructed figures and wrote the manuscript.

