## supplemental figures and table for "HMGB1/CXCL12 heterocomplex activates CXCR4 to suppress the ability of Müller glia to become proliferating progenitor-like cells in the chick retina"

**Supplemental Figure 1: Patterns of expression of *HMGB1, HMGB2, HMGB3* and *CXCR4* in embryonic chick retina.** These scRNA-libraries were generated and analyzed previously (Campbell et al., 2021)**.** scRNA-seq was used to identify patterns of expression of *HMGB1, HMGB2, HMGB3* and *CXCR4*-related factors in embryonic retinal cells. scRNA-seq libraries were generated from embryonic retinal cells at E5, E8, E12, and E15 (**a,b**). UMAP plots indicate cell types that were identified by expression of cell-distinguishing genes for progenitor cells (**c**) or mature glia (**d**), with cells expressing 2+ genes denoted in black. Expression patterns of *HMGB1, HMGB2, HMGB3* and *CXCR4* are illustrated in UMAP heatmaps (**c**). RPCs, iMG and mMG were subsetted are analyzed by sample identity (developmental timepoint). The dotplot in **f** illustrates expression levels and percent expression of genes. *$p < 0.0001$ difference compared to E8, E12 or E15 stages of development and #$p<0.001$ difference compared to E5 or E8 stages of development; Wilcoxon Rank Sum Test with Bonferroni correction. Abbreviations: iMG, immature Müller glia; mMG, mature Müller glia; RPC, retinal progenitor cell.

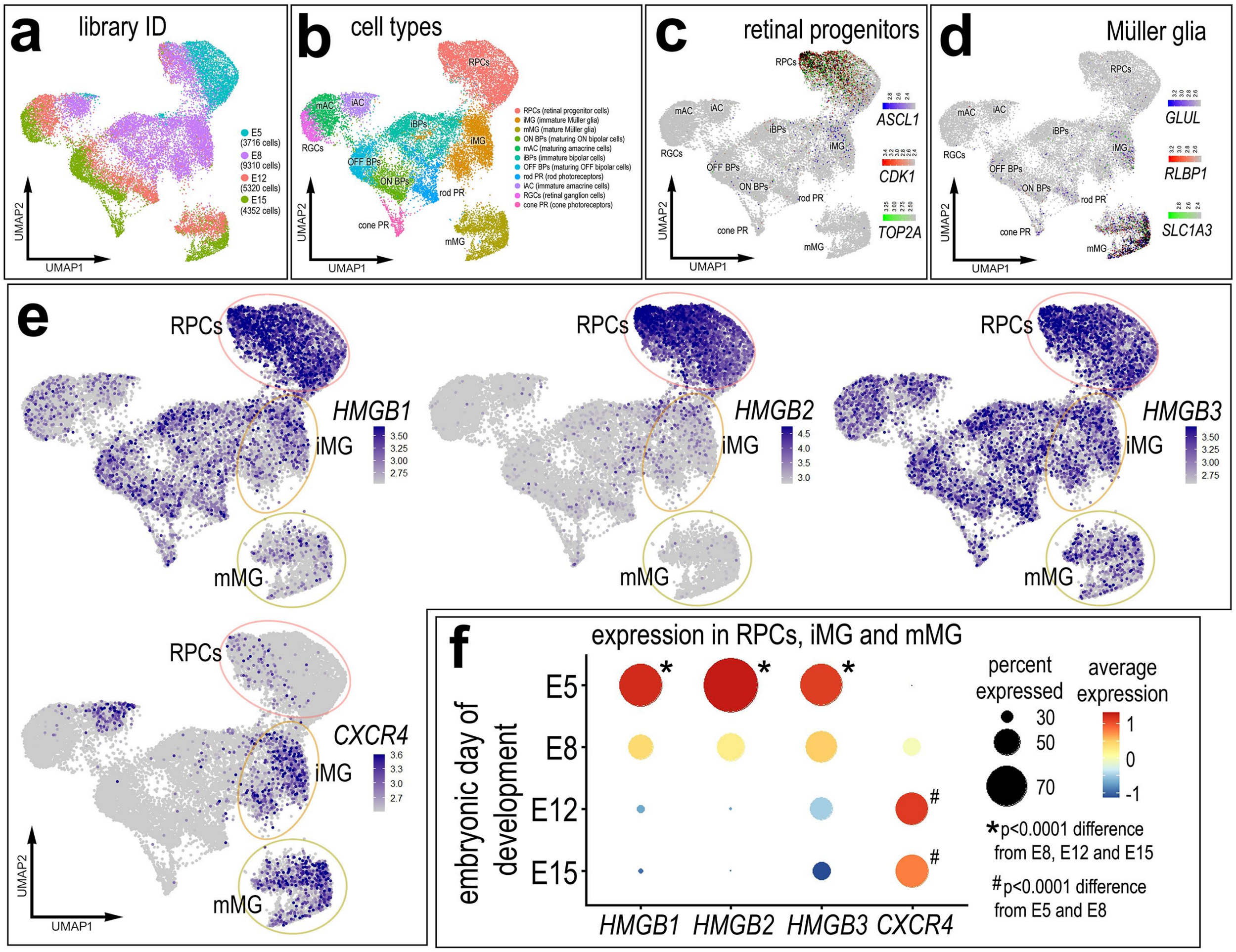

a
library ID
E5 (3716 cells)
E8 (9310 cells)
E12 (5320 cells)
E15 (4352 cells)
UMAP1
UMAP2
b
cell types
RPCs (retinal progenitor cells)
iMG (immature Müller glia)
mMG (mature Müller glia)
ON BPs (maturing ON bipolar cells)
mAC (maturing amacrine cells)
iBPs (immature bipolar cells)
OFF BPs (maturing OFF bipolar cells)
rod PR (rod photoreceptors)
iAC (immature amacrine cells)
RGCs (retinal ganglion cells)
cone PR (cone photoreceptors)
c
retinal progenitors
ASCL1
CDK1
TOP2A
d
Müller glia
GLUL
RLBP1
SLC1A3
e
RPCs
iMG
mMG
HMGB1
HMGB2
HMGB3
CXCR4
f
expression in RPCs, iMG and mMG
embryonic day of development
E5
E8
E12
E15
HMGB1
HMGB2
HMGB3
CXCR4
percent expressed
30
50
70
average expression
1
0
-1
*p<0.0001 difference from E8, E12 and E15
#p<0.0001 difference from E5 and E8

**Supplemental Figure 2: Immunolabeling for HMGB1 in different types of retinal neurons.** Sections of undamaged retinas were labeled with antibodies to HMGB1 (green) and Sox2 (red) or calretinin (red). Confocal microscopy was used to obtain single optical sections that were less than 2 μm thick. Hollow arrows (yellow) indicate DAPI-positive nuclei that are negative for HMGB1. Arrows (yellow) indicate DAPI-intense nuclei with HMGB1 exported out of the nuclei into the peri-nuclear cytoplasm. The calibration bars in **d** represent 50 μm and applies to **a-d**. Areas of the ONL, INL and GCL that are indicated by cyan rectangles are enlarged 2-fold in adjacent panels. Abbreviations: ONL – outer nuclear layer, INL – inner nuclear layer, IPL – inner plexiform layer, GCL – ganglion cell layer, ns – not significant. NMDA, N-methyl-D-aspartate.

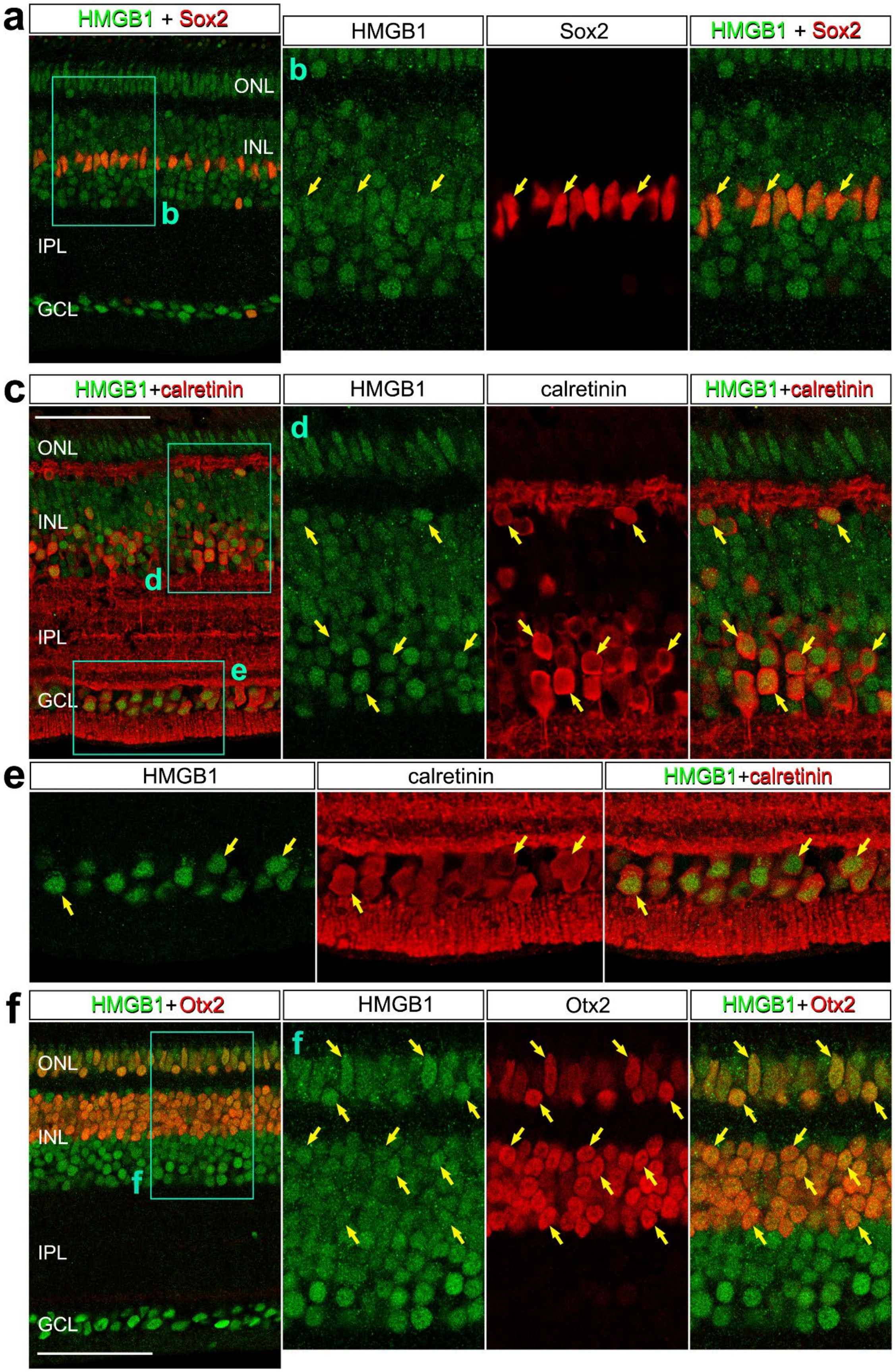

a
HMGB1 + Sox2
HMGB1
Sox2
HMGB1 + Sox2
ONL
INL
IPL
GCL
b
c
HMGB1+calretinin
HMGB1
calretinin
HMGB1+calretinin
d
e
HMGB1
calretinin
HMGB1+calretinin
f
HMGB1+Otx2
HMGB1
Otx2
HMGB1+Otx2

**Supplemental Figure 3: Otx2 remains confined to the nuclei of bipolar cells and photoreceptors in damaged retinas.** Sections of undamaged retinas were labeled with antibodies to HMGB1 (green) and Sox2 (red) or calretinin (red). Confocal microscopy was used to obtain single optical sections that were less than 2 µm thick. Hollow arrows (yellow) indicate DAPI-positive nuclei that are negative for HMGB1. Arrows (yellow) indicate DAPI-intense nuclei with HMGB1 exported out of the nuclei into the peri-nuclear cytoplasm. Solid arrow heads (yellow) indicate the nuclei of bipolar cells and photoreceptors labeled for DAPI, HMGB1 and Otx2. The calibration bar represents 50 µm and applies to unenlarged panels. Areas of the ON and INL and GCL are indicated by yellow boxes are enlarged 2-fold in adjacent panels.  Abbreviations: ONL – outer nuclear layer, INL – inner nuclear layer, IPL – inner plexiform layer, GCL – ganglion cell layer, ns – not significant. NMDA, N-methyl-D-aspartate.

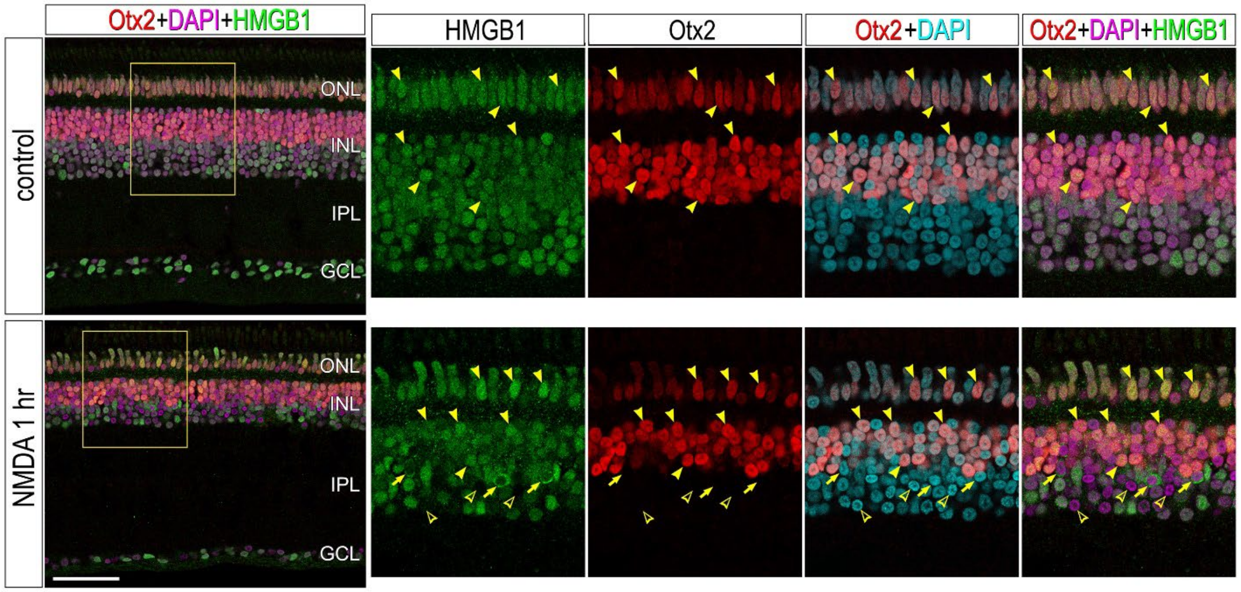

**Supplemental Figure 4: Inhibition of HMGB1 secretion, HMGB1/CXCL12 heterocomplex or CXCR4 has no effect on cell death or reactive microglia in damaged retinas.** Eyes were injected with NMDA+vehicle (control) or NMDA+inhibitor (treated) at P10, EdU+vehicle or inhibitor at P11 and P12, and retinas harvested at P13, 24 hrs after the last injection. Drugs included Lys2519024 (CXCR4 antagonist), Plerixafor (CXCR4 antagonist), inflachromene (inhibitor of HMGB1 secretion), and HPB08 (inhibitor of the HMGB1/CXCL12 heterocomplex). Vertical sections of the retina were labeled for fragmented DNA (TUNEL; a) or CD45 (red) and EdU (green; f). Arrows indicate the nuclei of microglia labeled for EdU and CD45. The calibration bar represents 50 µm (**a,f**). Histograms (**b-e**, **g-j**) illustrate the mean (bar ± SD), each dot represents one biological replicate, and blue lines connect counts from control and treated retinas from one individual. Significance of difference (p-values) was determined by using a paired t-test. Abbreviations: ONL – outer nuclear layer, INL – inner nuclear layer, IPL – inner plexiform layer, GCL – ganglion cell layer, ns – not significant, NMDA, N-methyl-D-aspartate, EdU- 5-Ethynyl-2-deoxyuridine.

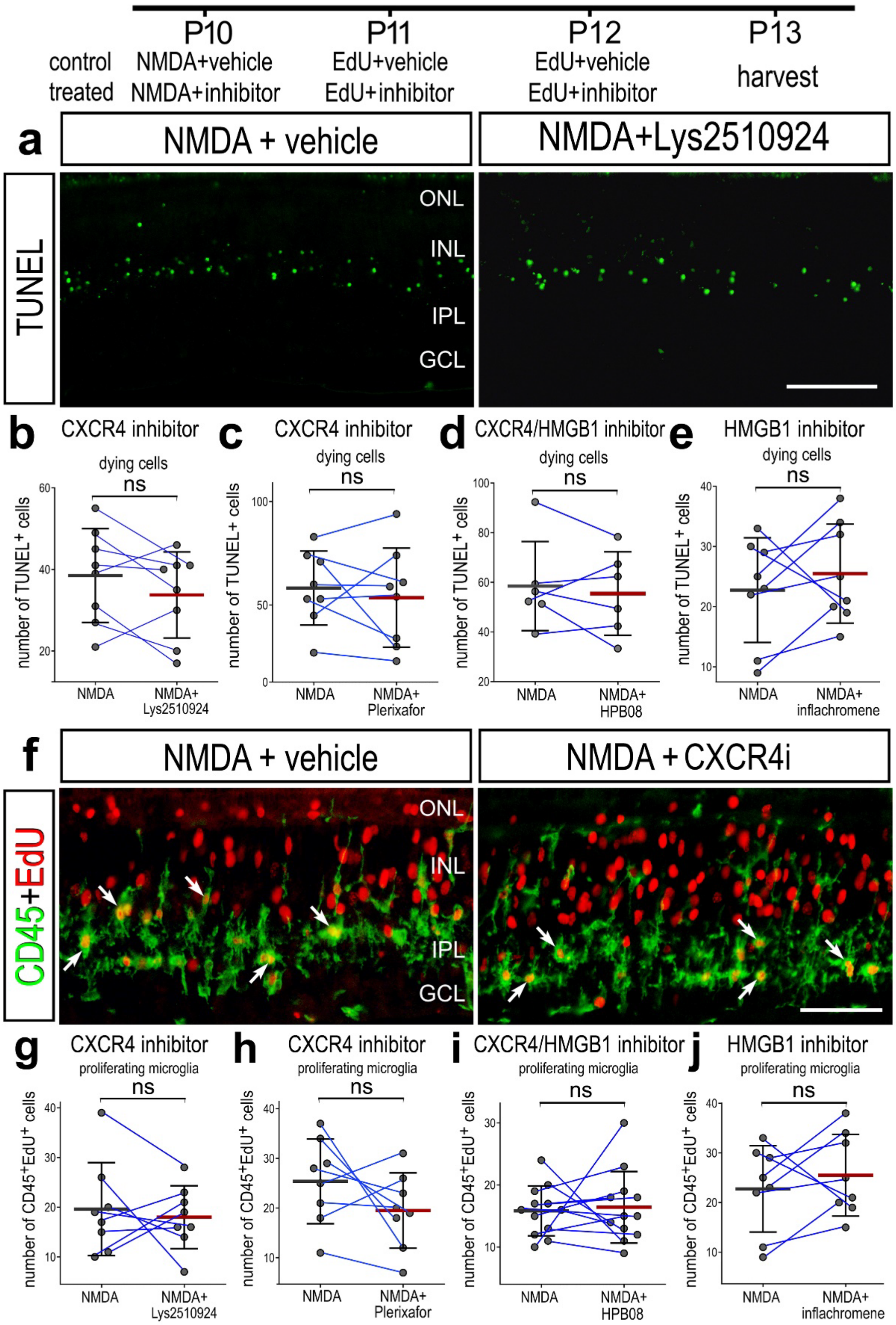

P10
P11
P12
P13
control
treated
NMDA+vehicle
NMDA+inhibitor
EdU+vehicle
EdU+inhibitor
EdU+vehicle
EdU+inhibitor
harvest
a
NMDA + vehicle
NMDA+Lys2510924
TUNEL
ONL
INL
IPL
GCL
b
CXCR4 inhibitor
dying cells
ns
number of TUNEL+ cells
NMDA
NMDA+ Lys2510924
c
CXCR4 inhibitor
dying cells
ns
number of TUNEL+ cells
NMDA
NMDA+ Plerixafor
d
CXCR4/HMGB1 inhibitor
dying cells
ns
number of TUNEL+ cells
NMDA
NMDA+ HPB08
e
HMGB1 inhibitor
dying cells
ns
number of TUNEL+ cells
NMDA
NMDA+ inflachromene
f
NMDA + vehicle
NMDA + CXCR4i
CD45+EdU
ONL
INL
IPL
GCL
g
CXCR4 inhibitor
proliferating microglia
ns
number of CD45+EdU+ cells
NMDA
NMDA+ Lys2510924
h
CXCR4 inhibitor
proliferating microglia
ns
number of CD45+EdU+ cells
NMDA
NMDA+ Plerixafor
i
CXCR4/HMGB1 inhibitor
proliferating microglia
ns
number of CD45+EdU+ cells
NMDA
NMDA+ HPB08
j
HMGB1 inhibitor
proliferating microglia
ns
number of CD45+EdU+ cells
NMDA
NMDA+ inflachromene

**Supplemental Figure 5: CXCR4 agonist has no effect on the proliferation of MGPCs in damaged retinas.** Eyes were injected with NMDA+vehicle (control) or NMDA+ATI-2341 (treated) at P10, EdU+ ATI-2341 or inhibitor at P11 and P12, and retinas harvested at P13, 24 hrs after the last injection. Vertical sections of the retina were labeled for EdU (red; **a**) and Sox2 (green; **a**). Arrows indicate the nuclei of the nuclei of proliferating MGPCs. The calibration bar represents 50 µm (**a**). The histogram in **b** illustrates the mean (bar ± SD), each dot represents one biological replicate, and blue lines connect counts from control and treated retinas from one individual. Significance of difference (p-values) was determined by using a paired t-test. Abbreviations: ONL – outer nuclear layer, INL – inner nuclear layer, IPL – inner plexiform layer, GCL – ganglion cell layer, ns – not significant, NMDA, N-methyl-D-aspartate, EdU- 5-Ethynyl-2-deoxyuridine.

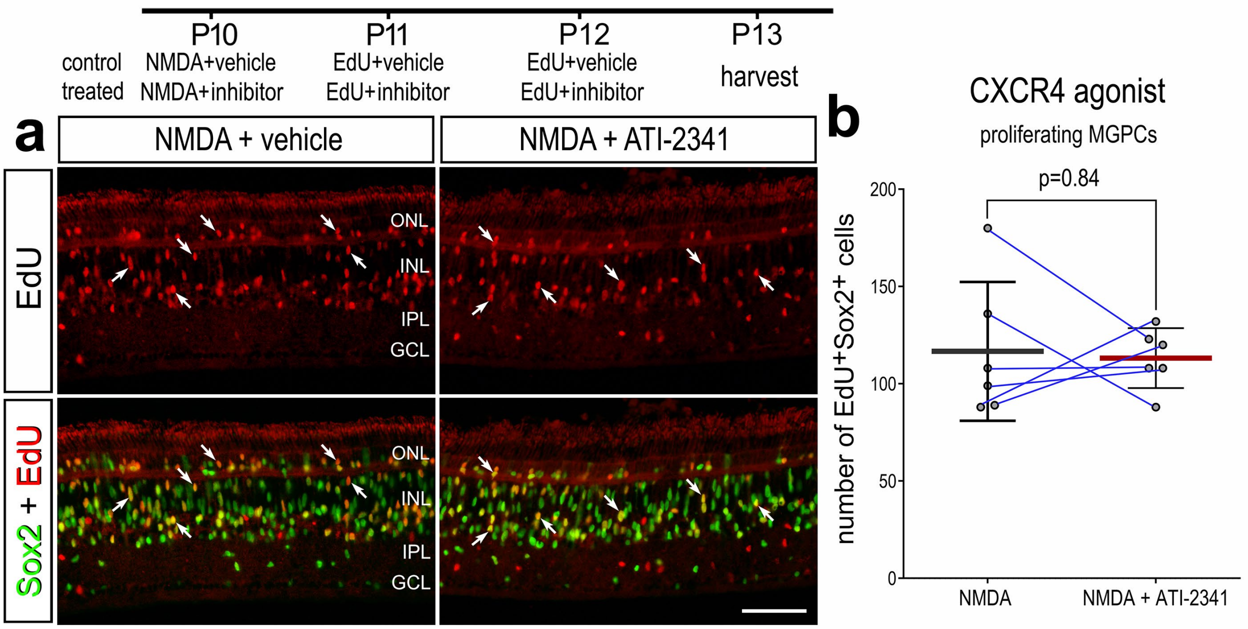
P10
P11
P12
P13
control treated
NMDA+vehicle
NMDA+inhibitor
EdU+vehicle
EdU+inhibitor
EdU+vehicle
EdU+inhibitor
harvest
a
NMDA + vehicle
NMDA + ATI-2341
EdU
Sox2 + EdU
ONL
INL
IPL
GCL
b
CXCR4 agonist
proliferating MGPCs
p=0.84
number of EdU+Sox2+ cells
200
150
100
50
0
NMDA
NMDA + ATI-2341

### Supplemental table: statistics for Dot plots and Violin plots

### Figure 2h

```
FindMarkers(NEW_MG_AGG, ident.1 = "resting MG", ident.2 = "return to resting
MG", features = c("HMGB1", "HMGB2", "HMGB3", "CXCR4"), min.pct = 0.001, logfc
.threshold = 0.001, test.use = "wilcox")
              p_val avg_log2FC pct.1 pct.2     p_val_adj
CXCR4 7.115201e-164  1.0289195 0.505 0.320 1.652505e-159
HMGB3  1.662374e-97  0.8355758 0.415 0.269  3.860865e-93
HMGB2  1.224076e-88  1.2729221 0.207 0.101  2.842917e-84
HMGB1  7.503030e-26  0.2903345 0.591 0.513  1.742579e-21
```

```
FindMarkers(NEW_MG_AGG, ident.1 = "resting MG", ident.2 = "returning farther
toward resting MG", features = c("HMGB1", "HMGB2", "HMGB3", "CXCR4"), min.pct
= 0.001, logfc.threshold = 0.001, test.use = "wilcox")
             p_val avg_log2FC pct.1 pct.2    p_val_adj
HMGB3 1.863497e-43  1.0514065 0.415 0.235 4.327972e-39
CXCR4 4.519415e-32  0.8152181 0.505 0.358 1.049634e-27
HMGB1 9.621187e-24  0.4121138 0.591 0.442 2.234521e-19
HMGB2 8.547304e-21  1.2027315 0.207 0.108 1.985111e-16
```

```
FindMarkers(NEW_MG_AGG, ident.1 = "resting MG", ident.2 = "24hr activated MG"
, features = c("HMGB1", "HMGB2", "HMGB3", "CXCR4"), min.pct = 0.001, logfc.th
reshold = 0.001, test.use = "wilcox")
             p_val avg_log2FC pct.1 pct.2    p_val_adj
HMGB3 1.308861e-16 0.09948907 0.415 0.586 3.039829e-12
HMGB1 1.253605e-15 0.43348041 0.591 0.695 2.911498e-11
CXCR4 1.438958e-06 0.15314158 0.505 0.671 3.341980e-02
HMGB2 3.467845e-01 0.64357605 0.207 0.244 1.000000e+00
```

```
FindMarkers(NEW_MG_AGG, ident.1 = "resting MG", ident.2 = "mix activated MG",
features = c("HMGB1", "HMGB2", "HMGB3", "CXCR4"), min.pct = 0.001, logfc.thre
shold = 0.001, test.use = "wilcox")
              p_val avg_log2FC pct.1 pct.2     p_val_adj
CXCR4 5.058427e-185 1.03027117 0.505 0.331 1.174820e-180
HMGB2  3.390765e-65 1.14499339 0.207 0.125  7.875052e-61
HMGB3  1.310402e-62 0.60666259 0.415 0.319  3.043409e-58
HMGB1  7.845575e-08 0.03008212 0.591 0.546  1.822135e-03
```

```
FindMarkers(NEW_MG_AGG, ident.1 = "resting MG", ident.2 = "2do FI", features
= c("HMGB1", "HMGB2", "HMGB3", "CXCR4"), min.pct = 0.001, logfc.threshold = 0
.001, test.use = "wilcox")
              p_val   avg_log2FC pct.1 pct.2     p_val_adj
CXCR4 1.458332e-201  1.319712384 0.505 0.282 3.386975e-197
HMGB1  3.882776e-21  0.295955827 0.591 0.540  9.017748e-17
HMGB3  1.232840e-08  0.258696063 0.415 0.378  2.863270e-04
HMGB2  6.091869e-02 -0.003255947 0.207 0.224  1.000000e+00
```

```
FindMarkers(NEW_MG_AGG, ident.1 = "resting MG", ident.2 = "FI de-differentiat
ing", features = c("HMGB1", "HMGB2", "HMGB3", "CXCR4"), min.pct = 0.001, logf
c.threshold = 0.001, test.use = "wilcox")
              p_val avg_log2FC pct.1 pct.2     p_val_adj
CXCR4 1.582595e-248 1.38513240 0.505 0.158 3.675578e-244
HMGB3 5.167024e-101 0.50447537 0.415 0.183  1.200041e-96
HMGB1  2.388169e-29 0.01152606 0.591 0.387  5.546522e-25
HMGB2  4.034925e-28 0.17825309 0.207 0.115  9.371112e-24


FindMarkers(NEW_MG_AGG, ident.1 = "resting MG", ident.2 = c("MGPC 1", "MGPC 2
", "MGPC 3"), features = c("HMGB1", "HMGB2", "HMGB3", "CXCR4"), min.pct = 0.0
01, logfc.threshold = 0.001, test.use = "wilcox")
              p_val avg_log2FC pct.1 pct.2     p_val_adj
HMGB2  0.000000e+00 -1.7356055 0.207 0.506  0.000000e+00
HMGB1  0.000000e+00 -1.6524209 0.591 0.857  0.000000e+00
HMGB3 1.233909e-128 -0.6225387 0.415 0.596 2.865755e-124
CXCR4 5.822971e-113  0.9293576 0.505 0.412 1.352385e-108


FindMarkers(NEW_MG_AGG, ident.1 = "returning farther toward resting MG", iden
t.2 = c("MGPC 1", "MGPC 2", "MGPC 3"), features = c("HMGB1", "HMGB2", "HMGB3"
, "CXCR4"), min.pct = 0.001, logfc.threshold = 0.001, test.use = "wilcox")
              p_val avg_log2FC pct.1 pct.2     p_val_adj
HMGB1  0.000000e+00 -2.0645347 0.442 0.857  0.000000e+00
HMGB2 2.709553e-200 -2.9383369 0.108 0.506 6.292936e-196
HMGB3 4.808884e-163 -1.6739452 0.235 0.596 1.116863e-158
CXCR4  2.125711e-01  0.1141395 0.358 0.412  1.000000e+00


FindMarkers(NEW_MG_AGG, ident.1 = "return to resting MG", ident.2 = c("MGPC 1
", "MGPC 2", "MGPC 3"), features = c("HMGB1", "HMGB2", "HMGB3", "CXCR4"), min
.pct = 0.001, logfc.threshold = 0.001, test.use = "wilcox")
             p_val  avg_log2FC pct.1 pct.2    p_val_adj
HMGB2 0.000000e+00 -3.00852757 0.101 0.506 0.000000e+00
HMGB1 0.000000e+00 -1.94275542 0.513 0.857 0.000000e+00
HMGB3 0.000000e+00 -1.45811450 0.269 0.596 0.000000e+00
CXCR4 4.380592e-27 -0.09956191 0.320 0.412 1.017393e-22


FindMarkers(NEW_MG_AGG, ident.1 = "24hr activated MG", ident.2 = c("MGPC 1",
"MGPC 2", "MGPC 3"), features = c("HMGB1", "HMGB2", "HMGB3", "CXCR4"), min.pc
t = 0.001, logfc.threshold = 0.001, test.use = "wilcox")
              p_val avg_log2FC pct.1 pct.2    p_val_adj
HMGB2  0.000000e+00 -2.3791815 0.244 0.506  0.00000e+00
CXCR4  0.000000e+00  0.7762160 0.671 0.412  0.00000e+00
HMGB1  0.000000e+00 -2.0859013 0.695 0.857  0.00000e+00
HMGB3 1.307272e-142 -0.7220278 0.586 0.596 3.03614e-138


FindMarkers(NEW_MG_AGG, ident.1 = "mix activated MG", ident.2 = c("MGPC 1", "
MGPC 2", "MGPC 3"), features = c("HMGB1", "HMGB2", "HMGB3", "CXCR4"), min.pct
= 0.001, logfc.threshold = 0.001, test.use = "wilcox")
            p_val avg_log2FC pct.1 pct.2    p_val_adj
```

```
HMGB2 0.00000e+00 -2.8805988 0.125 0.506 0.000000e+00
HMGB1 0.00000e+00 -1.6825030 0.546 0.857 0.000000e+00
HMGB3 0.00000e+00 -1.2292013 0.319 0.596 0.000000e+00
CXCR4 5.13163e-35 -0.1009136 0.331 0.412 1.191821e-30
```

```
FindMarkers(NEW_MG_AGG, ident.1 = "2do FI", ident.2 = c("MGPC 1", "MGPC 2", "
MGPC 3"), features = c("HMGB1", "HMGB2", "HMGB3", "CXCR4"), min.pct = 0.001,
logfc.threshold = 0.001, test.use = "wilcox")
              p_val avg_log2FC pct.1 pct.2     p_val_adj
HMGB1  0.000000e+00 -1.9483767 0.540 0.857  0.000000e+00
HMGB2  0.000000e+00 -1.7323495 0.224 0.506  0.000000e+00
HMGB3 9.580957e-248 -0.8812348 0.378 0.596 2.225177e-243
CXCR4  1.296025e-65 -0.3903548 0.282 0.412  3.010017e-61
```

```
FindMarkers(NEW_MG_AGG, ident.1 = "FI de-differentiating", ident.2 = c("MGPC
1", "MGPC 2", "MGPC 3"), features = c("HMGB1", "HMGB2", "HMGB3", "CXCR4"), mi
n.pct = 0.001, logfc.threshold = 0.001, test.use = "wilcox")
              p_val avg_log2FC pct.1 pct.2     p_val_adj
HMGB1  0.000000e+00 -1.6639470 0.387 0.857  0.000000e+00
HMGB3  0.000000e+00 -1.1270141 0.183 0.596  0.000000e+00
HMGB2  0.000000e+00 -1.9138585 0.115 0.506  0.000000e+00
CXCR4 6.628534e-154 -0.4557748 0.158 0.412 1.539477e-149
```

### Figure 3d

### Stats chick early MG HMGB1 CXCR4

```
FindMarkers(MullerG, ident.1 = "saline", ident.2 = "NMDA 3hr", group.by =
"orig.ident", features = c("HMGB1", "HMGB2", "CXCR4"), min.pct = 0.001,
logfc.threshold = 0.001, test.use = "wilcox")
              p_val avg_log2FC pct.1 pct.2    p_val_adj
CXCR4 2.348800e-57 -1.0631684 0.569 0.804 4.377693e-53
HMGB2 8.419563e-19  0.7648187 0.556 0.446 1.569238e-14
HMGB1 7.461235e-13  0.5371078 0.765 0.826 1.390625e-08
```

```
FindMarkers(MullerG, ident.1 = "saline", ident.2 = "NMDA 12hr", group.by =
"orig.ident", features = c("HMGB1", "HMGB2", "CXCR4"), min.pct = 0.001,
logfc.threshold = 0.001, test.use = "wilcox")
              p_val avg_log2FC pct.1 pct.2    p_val_adj
CXCR4 5.340701e-49 -0.4676446 0.569 0.816 9.953999e-45
HMGB2 2.833513e-20  0.7090112 0.556 0.496 5.281102e-16
HMGB1 3.692464e-07  0.3535070 0.765 0.845 6.882014e-03
```

```
FindMarkers(MullerG, ident.1 = "saline", ident.2 = "NMDA 48hr", group.by =
"orig.ident", features = c("HMGB1", "HMGB2", "CXCR4"), min.pct = 0.001,
logfc.threshold = 0.001, test.use = "wilcox")
              p_val avg_log2FC pct.1 pct.2    p_val_adj
HMGB1 1.238416e-78 -1.0388575 0.765 0.850 2.308159e-74
HMGB2 1.650906e-19 -0.5393029 0.556 0.686 3.076958e-15
CXCR4 2.042671e-07  0.5354853 0.569 0.579 3.807130e-03
```

```
FindMarkers(MullerG, ident.1 = "NMDA 3hr", ident.2 = "NMDA 48hr", group.by =
"orig.ident", features = c("HMGB1", "HMGB2", "CXCR4"), min.pct = 0.001,
logfc.threshold = 0.001, test.use = "wilcox")
              p_val avg_log2FC pct.1 pct.2    p_val_adj
HMGB1 3.314960e-87  -1.575965 0.826 0.850 6.178422e-83
CXCR4 3.890728e-77   1.598654 0.804 0.579 7.251539e-73
HMGB2 4.993352e-49  -1.304122 0.446 0.686 9.306610e-45
```

```
FindMarkers(MullerG, ident.1 = "NMDA 3hr", ident.2 = "NMDA 12hr", group.by =
"orig.ident", features = c("HMGB1", "HMGB2", "CXCR4"), min.pct = 0.001,
logfc.threshold = 0.001, test.use = "wilcox")
              p_val avg_log2FC pct.1 pct.2    p_val_adj
CXCR4 8.529116e-13  0.59552376 0.804 0.816 1.589657e-08
HMGB1 4.748511e-05 -0.18360076 0.826 0.845 8.850275e-01
HMGB2 2.073796e-02 -0.05580751 0.446 0.496 1.000000e+00
```

```
FindMarkers(MullerG, ident.1 = "NMDA 48hr", ident.2 = "NMDA 12hr", group.by =
"orig.ident", features = c("HMGB1", "HMGB2", "CXCR4"), min.pct = 0.001,
logfc.threshold = 0.001, test.use = "wilcox")
              p_val avg_log2FC pct.1 pct.2    p_val_adj
HMGB1 4.135223e-109   1.392365 0.850 0.845 7.707229e-105
CXCR4  7.723067e-86  -1.003130 0.579 0.816  1.439425e-81
HMGB2  8.321024e-61   1.248314 0.686 0.496  1.550872e-56
```

### Figure 3e

Stats for Dot plots chick early NMDA aggregate

#### Whole retina

```
FindMarkers(early_AGG, ident.1 = "saline", ident.2 = "NMDA 3hr", group.by = "
orig.ident", features = c("HMGB1", "HMGB2", "HMGB3"), min.pct = 0.001, logfc.
threshold = 0.001, test.use = "wilcox")
              p_val avg_log2FC pct.1 pct.2    p_val_adj
HMGB1 8.044307e-101  0.3755542 0.814 0.722 1.499298e-96
HMGB3  9.783390e-82  0.4503501 0.561 0.446 1.823428e-77
HMGB2  5.751415e-64  0.3757570 0.538 0.406 1.071949e-59
```

```
FindMarkers(early_AGG, ident.1 = "saline", ident.2 = "NMDA 12hr", group.by =
"orig.ident", features = c("HMGB1", "HMGB2", "HMGB3"), min.pct = 0.001, logfc
.threshold = 0.001, test.use = "wilcox")
              p_val avg_log2FC pct.1 pct.2     p_val_adj
HMGB2 6.091197e-107 0.43568928 0.538 0.407 1.135277e-102
HMGB3  1.117732e-34 0.22189493 0.561 0.493  2.083228e-30
HMGB1  1.536862e-05 0.05396064 0.814 0.791  2.864404e-01
```

```
FindMarkers(early_AGG, ident.1 = "saline", ident.2 = "NMDA 48hr", group.by =
"orig.ident", features = c("HMGB1", "HMGB2", "HMGB3"), min.pct = 0.001, logfc
.threshold = 0.001, test.use = "wilcox")
             p_val   avg_log2FC pct.1 pct.2    p_val_adj
HMGB1 3.771705e-25 -0.159163217 0.814 0.832 7.029704e-21
HMGB2 9.600227e-15 -0.166167765 0.538 0.576 1.789290e-10
HMGB3 9.895963e-01 -0.006959796 0.561 0.563 1.000000e+00
```

```
FindMarkers(early_AGG, ident.1 = "NMDA 3hr", ident.2 = "NMDA 12hr", group.by
= "orig.ident", features = c("HMGB1", "HMGB2", "HMGB3"), min.pct = 0.001, log
fc.threshold = 0.001, test.use = "wilcox")
```

```
             p_val  avg_log2FC pct.1 pct.2    p_val_adj
HMGB1 3.576756e-74 -0.32159355 0.722 0.791 6.666357e-70
HMGB3 2.508601e-21 -0.22845515 0.446 0.493 4.675531e-17
HMGB2 6.841617e-01  0.05993228 0.406 0.407 1.000000e+00
```

```
FindMarkers(early_AGG, ident.1 = "NMDA 3hr", ident.2 = "NMDA 48hr", group.by
= "orig.ident", features = c("HMGB1", "HMGB2", "HMGB3"), min.pct = 0.001, log
fc.threshold = 0.001, test.use = "wilcox")
              p_val avg_log2FC pct.1 pct.2     p_val_adj
HMGB1 4.662431e-183 -0.5347174 0.722 0.832 8.689840e-179
HMGB2 3.888557e-112 -0.5419248 0.406 0.576 7.247493e-108
HMGB3  9.379865e-78 -0.4573099 0.446 0.563  1.748219e-73
```

```
FindMarkers(early_AGG, ident.1 = "NMDA 12hr", ident.2 = "NMDA 48hr", group.by
= "orig.ident", features = c("HMGB1", "HMGB2", "HMGB3"), min.pct = 0.001, log
fc.threshold = 0.001, test.use = "wilcox")
              p_val avg_log2FC pct.1 pct.2     p_val_adj
HMGB2 5.415493e-183 -0.6018570 0.407 0.576 1.009340e-178
HMGB1  1.973790e-49 -0.2131239 0.791 0.832  3.678751e-45
HMGB3  2.131339e-32 -0.2288547 0.493 0.563  3.972389e-28
```

### Inner retinal neurons – HCs+BPs+Acs+RGCs

```
FindMarkers(inners, ident.1 = "saline", ident.2 = "NMDA 3hr", group.by = "ori
g.ident", features = c("HMGB1", "HMGB2", "HMGB3"), min.pct = 0.001, logfc.thr
eshold = 0.001, test.use = "wilcox")
             p_val avg_log2FC pct.1 pct.2    p_val_adj
HMGB1 3.641263e-86  0.3830799 0.833 0.708 6.786586e-82
HMGB3 2.166150e-64  0.4615151 0.553 0.398 4.037271e-60
HMGB2 1.037889e-38  0.3119302 0.490 0.347 1.934417e-34
```

```
FindMarkers(inners, ident.1 = "saline", ident.2 = "NMDA 12hr", group.by = "or
ig.ident", features = c("HMGB1", "HMGB2", "HMGB3"), min.pct = 0.001, logfc.th
reshold = 0.001, test.use = "wilcox")
             p_val avg_log2FC pct.1 pct.2    p_val_adj
HMGB2 5.612042e-96 0.47885143 0.490 0.316 1.045972e-91
HMGB3 9.703381e-44 0.26738145 0.553 0.433 1.808516e-39
HMGB1 9.658805e-06 0.04384278 0.833 0.783 1.800208e-01
```

```
FindMarkers(inners, ident.1 = "saline", ident.2 = "NMDA 48hr", group.by = "or
ig.ident", features = c("HMGB1", "HMGB2", "HMGB3"), min.pct = 0.001, logfc.th
reshold = 0.001, test.use = "wilcox")
            p_val  avg_log2FC pct.1 pct.2 p_val_adj
HMGB2 0.004393737 -0.09779388 0.490 0.507         1
HMGB3 0.012003404  0.03906946 0.553 0.537         1
HMGB1 0.390572187 -0.03066336 0.833 0.823         1
```

```
FindMarkers(inners, ident.1 = "NMDA 3hr", ident.2 = "NMDA 12hr", group.by = "
orig.ident", features = c("HMGB1", "HMGB2", "HMGB3"), min.pct = 0.001, logfc.
threshold = 0.001, test.use = "wilcox")
             p_val avg_log2FC pct.1 pct.2    p_val_adj
HMGB1 1.078172e-55 -0.3392371 0.708 0.783 2.009497e-51
HMGB3 7.696652e-08 -0.1941336 0.398 0.433 1.434502e-03
HMGB2 1.821763e-04  0.1669212 0.347 0.316 1.000000e+00
```

```
FindMarkers(inners, ident.1 = "NMDA 3hr", ident.2 = "NMDA 48hr", group.by = "
orig.ident", features = c("HMGB1", "HMGB2", "HMGB3"), min.pct = 0.001, logfc.
threshold = 0.001, test.use = "wilcox")
             p_val avg_log2FC pct.1 pct.2    p_val_adj
HMGB1 2.690792e-84 -0.4137433 0.708 0.823 5.015098e-80
HMGB2 7.030127e-50 -0.4097241 0.347 0.507 1.310275e-45
HMGB3 1.612203e-47 -0.4224456 0.398 0.537 3.004824e-43
```

```
FindMarkers(inners, ident.1 = "NMDA 12hr", ident.2 = "NMDA 48hr", group.by =
"orig.ident", features = c("HMGB1", "HMGB2", "HMGB3"), min.pct = 0.001, logfc
.threshold = 0.001, test.use = "wilcox")
              p_val  avg_log2FC pct.1 pct.2     p_val_adj
HMGB2 4.094822e-115 -0.57664531 0.316 0.507 7.631929e-111
HMGB3  1.881264e-28 -0.22831199 0.433 0.537  3.506300e-24
HMGB1  4.324209e-07 -0.07450614 0.783 0.823  8.059461e-03
```

### Microglia

```
FindMarkers(microglia, ident.1 = "saline", ident.2 = "NMDA 3hr", group.by = "
orig.ident", features = c("HMGB1", "HMGB2", "HMGB3", "S100A4", "S100A6", "S10
0A11"), min.pct = 0.001, logfc.threshold = 0.001, test.use = "wilcox")
               p_val avg_log2FC pct.1 pct.2    p_val_adj
HMGB1   5.717883e-12  1.6759770 0.588 0.225 1.065699e-07
S100A11 3.499773e-06  1.1617115 0.286 0.063 6.522876e-02
HMGB2   7.816230e-05  1.2237588 0.196 0.036 1.000000e+00
HMGB3   7.148443e-04  0.9793053 0.186 0.054 1.000000e+00
S100A6  4.267694e-03 -0.8226726 0.611 0.667 1.000000e+00
S100A4  6.368093e-01 -0.1306969 0.643 0.577 1.000000e+00
```

```
FindMarkers(microglia, ident.1 = "saline", ident.2 = "NMDA 12hr", group.by =
"orig.ident", features = c("HMGB1", "HMGB2", "HMGB3", "S100A4", "S100A6", "S1
00A11"), min.pct = 0.001, logfc.threshold = 0.001, test.use = "wilcox")
               p_val  avg_log2FC pct.1 pct.2    p_val_adj
S100A6  2.635980e-16 -0.88496746 0.611 0.846 4.912939e-12
HMGB1   5.039228e-11  1.04890960 0.588 0.422 9.392113e-07
S100A11 8.368583e-03  0.51892836 0.286 0.212 1.000000e+00
HMGB2   2.874254e-02  0.69090217 0.196 0.143 1.000000e+00
HMGB3   1.387874e-01  0.29155116 0.186 0.149 1.000000e+00
S100A4  1.759864e-01 -0.06747719 0.643 0.761 1.000000e+00
```

```
FindMarkers(microglia, ident.1 = "saline", ident.2 = "NMDA 48hr", group.by =
"orig.ident", features = c("HMGB1", "HMGB2", "HMGB3", "S100A4", "S100A6", "S1
00A11"), min.pct = 0.001, logfc.threshold = 0.001, test.use = "wilcox")
               p_val  avg_log2FC pct.1 pct.2    p_val_adj
S100A6  1.083494e-18 -0.78333732 0.611 0.947 2.019416e-14
HMGB1   1.390304e-12 -0.57994474 0.588 0.922 2.591248e-08
S100A11 1.662574e-12 -0.62489677 0.286 0.653 3.098706e-08
HMGB2   2.341187e-07 -0.18904134 0.196 0.445 4.363504e-03
HMGB3   1.940066e-06 -0.09768829 0.186 0.416 3.615895e-02
S100A4  3.033647e-04 -0.19699958 0.643 0.922 1.000000e+00
```

```
FindMarkers(microglia, ident.1 = "NMDA 3hr", ident.2 = "NMDA 12hr", group.by
= "orig.ident", features = c("HMGB1", "HMGB2", "HMGB3", "S100A4", "S100A6", "
S100A11"), min.pct = 0.001, logfc.threshold = 0.001, test.use = "wilcox")
               p_val  avg_log2FC pct.1 pct.2 p_val_adj
HMGB1   0.0005707851 -0.62706739 0.225 0.422         1
S100A11 0.0006818369 -0.64278314 0.063 0.212         1
HMGB2   0.0028767460 -0.53285665 0.036 0.143         1
```

```
HMGB3   0.0084273806 -0.68775418 0.054 0.149         1
S100A6  0.0236943019 -0.06229483 0.667 0.846         1
S100A4  0.1430900106  0.06321966 0.577 0.761         1


FindMarkers(microglia, ident.1 = "NMDA 3hr", ident.2 = "NMDA 48hr", group.by
= "orig.ident", features = c("HMGB1", "HMGB2", "HMGB3", "S100A4", "S100A6", "
S100A11"), min.pct = 0.001, logfc.threshold = 0.001, test.use = "wilcox")
              p_val  avg_log2FC pct.1 pct.2    p_val_adj
HMGB1   1.318847e-32 -2.25592173 0.225 0.922 2.458068e-28
S100A11 1.970096e-20 -1.78660826 0.063 0.653 3.671865e-16
HMGB2   3.324148e-13 -1.41280015 0.036 0.445 6.195547e-09
HMGB3   3.869224e-11 -1.07699363 0.054 0.416 7.211460e-07
S100A4  1.426371e-03 -0.06630273 0.577 0.922 1.000000e+00
S100A6  4.988801e-03  0.03933532 0.667 0.947 1.000000e+00


FindMarkers(microglia, ident.1 = "NMDA 12hr", ident.2 = "NMDA 48hr", group.by
= "orig.ident", features = c("HMGB1", "HMGB2", "HMGB3", "S100A4", "S100A6", "
S100A11"), min.pct = 0.001, logfc.threshold = 0.001, test.use = "wilcox")
              p_val avg_log2FC pct.1 pct.2    p_val_adj
HMGB1   2.884932e-48 -1.6288543 0.422 0.922 5.376937e-44
S100A11 5.589752e-26 -1.1438251 0.212 0.653 1.041818e-21
HMGB2   1.156794e-15 -0.8799435 0.143 0.445 2.156032e-11
HMGB3   1.551598e-11 -0.3892395 0.149 0.416 2.891869e-07
S100A4  7.256039e-03 -0.1295224 0.761 0.922 1.000000e+00
S100A6  6.608645e-01  0.1016301 0.846 0.947 1.000000e+00


FindMarkers(microglia, ident.1 = "saline", ident.2 = "NMDA 3hr", group.by = "
orig.ident", features = c("HMGB1", "HMGB2", "TLR4", "ENSGALG00000009239"), mi
n.pct = 0.01, logfc.threshold = 0.01, test.use = "wilcox")
                          p_val  avg_log2FC pct.1 pct.2    p_val_adj
HMGB1              3.318973e-22  2.00679101 0.588 0.225 6.185902e-18
HMGB2              5.417604e-06  2.49713902 0.196 0.036 1.009733e-01
TLR4               6.789750e-03  0.05190011 0.222 0.162 1.000000e+00
ENSGALG00000009239 7.813884e-01 -0.12947932 0.023 0.018 1.000000e+00


FindMarkers(microglia, ident.1 = "saline", ident.2 = "NMDA 12hr", group.by =
"orig.ident", features = c("HMGB1", "HMGB2", "TLR4", "ENSGALG00000009239"), m
in.pct = 0.01, logfc.threshold = 0.01, test.use = "wilcox")
  |++++++++++++++++++++++++++++++++++++++++++++++++++| 100% elapsed=00s
                          p_val avg_log2FC pct.1 pct.2    p_val_adj
HMGB1              7.828208e-29   1.202095 0.588 0.422 1.459021e-24
TLR4               3.348392e-06   1.559636 0.222 0.122 6.240732e-02
HMGB2              1.374435e-05   1.150579 0.196 0.143 2.561672e-01
ENSGALG00000009239 1.150974e-02  -1.040994 0.023 0.058 1.000000e+00


> FindMarkers(microglia, ident.1 = "saline", ident.2 = "NMDA 48hr", group.by
= "orig.ident", features = c("HMGB1", "HMGB2", "TLR4", "ENSGALG00000009239"),
min.pct = 0.01, logfc.threshold = 0.01, test.use = "wilcox")
                           p_val avg_log2FC pct.1 pct.2     p_val_adj
HMGB1              6.522306e-170 -0.6242892 0.588 0.922 1.215627e-165
HMGB2               7.278797e-19 -0.2660199 0.196 0.445  1.356622e-14
TLR4                4.195192e-06  0.5309703 0.222 0.343  7.819000e-02
ENSGALG00000009239  3.349167e-03 -0.8732268 0.023 0.065  1.000000e+00
```

**Retinal Ganglion cells – stressed but not dying**

```
FindMarkers(RGCs, ident.1 = "saline", ident.2 = "NMDA 3hr", group.by = "orig.
ident", features = c("HMGB1", "HMGB2"), min.pct = 0.01, logfc.threshold = 0.0
1, test.use = "poisson")
              p_val avg_log2FC pct.1 pct.2     p_val_adj
HMGB1 2.313825e-146   1.198673 0.990 0.928 4.312508e-142
HMGB2  4.519666e-30   1.461150 0.742 0.353  8.423754e-26
```

```
FindMarkers(RGCs, ident.1 = "saline", ident.2 = "NMDA 12hr", group.by = "orig
.ident", features = c("HMGB1", "HMGB2"), min.pct = 0.01, logfc.threshold = 0.
01, test.use = "poisson")
             p_val avg_log2FC pct.1 pct.2    p_val_adj
HMGB1 1.565338e-94  0.2142954 0.990 0.972 2.917477e-90
HMGB2 3.432716e-28  0.5201632 0.742 0.474 6.397896e-24
```

```
FindMarkers(RGCs, ident.1 = "saline", ident.2 = "NMDA 48hr", group.by = "orig
.ident", features = c("HMGB1", "HMGB2"), min.pct = 0.01, logfc.threshold = 0.
01, test.use = "poisson")
             p_val avg_log2FC pct.1 pct.2    p_val_adj
HMGB1 1.496008e-19 0.03597339 0.990 0.994 2.788259e-15
HMGB2 3.629435e-04 0.03271763 0.742 0.627 1.000000e+00
```

### Figure 10k

```
FindMarkers(insulin_FGF2_tr_MG, ident.1 = "resting MG", ident.2 = "MG 2do I+F
", features = c("HMGB1", "HMGB2", "HMGB3", "CXCR4"), min.pct = 0.001, logfc.t
hreshold = 0.001, test.use = "wilcox")
             p_val avg_log2FC pct.1 pct.2    p_val_adj
CXCR4 0.000000e+00  1.8473706 0.612 0.265 0.000000e+00
HMGB1 3.493917e-46  0.4670519 0.576 0.473 5.191612e-42
HMGB3 8.420468e-38  0.4945068 0.466 0.364 1.251197e-33
HMGB2 1.010304e-02  0.1357588 0.228 0.213 1.000000e+00
```

```
FindMarkers(insulin_FGF2_tr_MG, ident.1 = "resting MG", ident.2 = "MG 3do I+F
", features = c("HMGB1", "HMGB2", "HMGB3", "CXCR4"), min.pct = 0.001, logfc.t
hreshold = 0.001, test.use = "wilcox")
              p_val avg_log2FC pct.1 pct.2    p_val_adj
CXCR4 1.152903e-102  0.9764188 0.612 0.392 1.713099e-98
HMGB3  2.876388e-54  0.7209778 0.466 0.301 4.274024e-50
HMGB2  5.781485e-37  0.9710843 0.228 0.119 8.590709e-33
HMGB1  4.017795e-36  0.4167903 0.576 0.434 5.970041e-32
```

```
FindMarkers(insulin_FGF2_tr_MG, ident.1 = "resting MG", ident.2 = "MGPCs", fe
atures = c("HMGB1", "HMGB2", "HMGB3", "CXCR4"), min.pct = 0.001, logfc.thresh
old = 0.001, test.use = "wilcox")
              p_val avg_log2FC pct.1 pct.2     p_val_adj
HMGB1 5.387035e-227 -1.2607377 0.576 0.826 8.004595e-223
CXCR4  3.590123e-88  1.0583624 0.612 0.416  5.334564e-84
HMGB2  1.118937e-53 -1.0641779 0.228 0.389  1.662628e-49
HMGB3  3.267081e-03 -0.1061162 0.466 0.502  1.000000e+00
```

```
FindMarkers(insulin_FGF2_tr_MG, ident.1 = "MGPCs", ident.2 = "MG 2do I+F", fe
atures = c("HMGB1", "HMGB2", "HMGB3", "CXCR4"), min.pct = 0.001, logfc.thresh
old = 0.001, test.use = "wilcox")
             p_val avg_log2FC pct.1 pct.2    p_val_adj
HMGB1 0.000000e+00  1.7277896 0.826 0.473 0.000000e+00
HMGB2 4.545480e-83  1.1999366 0.389 0.213 6.754129e-79
CXCR4 2.689898e-53  0.7890082 0.416 0.265 3.996920e-49
HMGB3 1.076266e-43  0.6006230 0.502 0.364 1.599224e-39
```

```
FindMarkers(insulin_FGF2_tr_MG, ident.1 = "MGPCs", ident.2 = "MG 3do I+F", fe
atures = c("HMGB1", "HMGB2", "HMGB3", "CXCR4"), min.pct = 0.001, logfc.thresh
old = 0.001, test.use = "wilcox")
              p_val avg_log2FC pct.1 pct.2     p_val_adj
HMGB1 1.291571e-316  1.6775279 0.826 0.434 1.919145e-312
HMGB2 1.210869e-140  2.0352622 0.389 0.119 1.799230e-136
HMGB3  2.881085e-60  0.8270941 0.502 0.301  4.281005e-56
CXCR4  8.384556e-01 -0.0819436 0.416 0.392  1.000000e+00
```

### Figure 11f

```
FindMarkers(clod_tr_MullerGlia, ident.1 = "SS", ident.2 = "clod_SS", features
= c("HMGB1", "HMGB2", "HMGB3", "CXCR4"), group.by = "orig.ident", min.pct = 0
.001, logfc.threshold = 0.001, test.use = "wilcox")
             p_val  avg_log2FC pct.1 pct.2    p_val_adj
CXCR4 4.861778e-27  0.99251172 0.625 0.379 1.024182e-22
HMGB1 4.341971e-02  0.05231451 0.638 0.560 1.000000e+00
HMGB3 4.561004e-02  0.11427609 0.440 0.383 1.000000e+00
HMGB2 4.613230e-01 -0.24653044 0.223 0.235 1.000000e+00
```

```
FindMarkers(clod_tr_MullerGlia, ident.1 = "SS", ident.2 = "NM", features = c(
"HMGB1", "HMGB2", "HMGB3", "CXCR4"), group.by = "orig.ident", min.pct = 0.001
, logfc.threshold = 0.001, test.use = "wilcox")
             p_val avg_log2FC pct.1 pct.2    p_val_adj
HMGB1 2.439181e-08 0.41551661 0.638 0.645 0.0005138379
HMGB2 1.840988e-01 0.55210623 0.223 0.223 1.0000000000
HMGB3 5.333055e-01 0.27426431 0.440 0.513 1.0000000000
CXCR4 9.344853e-01 0.05469038 0.625 0.681 1.0000000000
```

```
FindMarkers(clod_tr_MullerGlia, ident.1 = "SS", ident.2 = "clod_NM", features
= c("HMGB1", "HMGB2", "HMGB3", "CXCR4"), group.by = "orig.ident", min.pct = 0
.001, logfc.threshold = 0.001, test.use = "wilcox")
             p_val avg_log2FC pct.1 pct.2    p_val_adj
CXCR4 6.028193e-26  1.1529884 0.625 0.425 1.269899e-21
HMGB3 1.062586e-01  0.3520667 0.440 0.462 1.000000e+00
HMGB1 2.942533e-01 -0.1670968 0.638 0.666 1.000000e+00
HMGB2 8.337225e-01  0.1730300 0.223 0.228 1.000000e+00
```

```
FindMarkers(clod_tr_MullerGlia, ident.1 = "NM", ident.2 = "clod_NM", features
= c("HMGB1", "HMGB2", "HMGB3", "CXCR4"), group.by = "orig.ident", min.pct = 0
.001, logfc.threshold = 0.001, test.use = "wilcox")
             p_val  avg_log2FC pct.1 pct.2    p_val_adj
CXCR4 9.107548e-29  1.09829801 0.681 0.425 1.918596e-24
HMGB1 1.284211e-08 -0.58261343 0.645 0.666 2.705318e-04
HMGB3 3.717393e-01  0.07780241 0.513 0.462 1.000000e+00
HMGB2 3.819900e-01 -0.37907624 0.223 0.228 1.000000e+00
```
